# Spectrally specific temporal analyses of spike-train responses to complex sounds: A unifying framework

**DOI:** 10.1101/2020.07.17.208330

**Authors:** Satyabrata Parida, Hari Bharadwaj, Michael G. Heinz

## Abstract

Significant scientific and translational questions remain in auditory neuroscience surrounding the neural correlates of perception. Relating perceptual and neural data collected from humans can be useful; however, human-based neural data are typically limited to evoked far-field responses, which lack anatomical and physiological specificity. Laboratory-controlled preclinical animal models offer the advantage of comparing single-unit and evoked responses from the same animals. This ability provides opportunities to develop invaluable insight into proper interpretations of evoked responses, which benefits both basic-science studies of neural mechanisms and translational applications, e.g., diagnostic development. However, these comparisons have been limited by a disconnect between the types of spectrotemporal analyses used with single-unit spike trains and evoked responses, which results because these response types are fundamentally different (point-process versus continuous-valued signals) even though the responses themselves are related. Here, we describe a unifying framework to study temporal coding of complex sounds that allows spike-train and evoked-response data to be analyzed and compared using the same advanced signal-processing techniques. The framework uses alternating-polarity peristimulus-time histograms computed from single-unit spike trains to allow advanced spectral analyses of both slow (envelope) and rapid (temporal fine structure) response components. Demonstrated benefits include: (1) novel spectrally specific temporal-coding measures that are less corrupted by analysis distortions due to hair-cell transduction, synaptic rectification, and neural stochasticity compared to previous metrics, e.g., the correlogram peak-height, (2) spectrally specific analyses of spike-train modulation coding (magnitude and phase), which can be directly compared to modern perceptually based models of speech intelligibility (e.g., that depend on modulation filter banks), and (3) superior spectral resolution in analyzing the neural representation of nonstationary sounds, such as speech and music. This unifying framework significantly expands the potential of preclinical animal models to advance our understanding of the physiological correlates of perceptual deficits in real-world listening following sensorineural hearing loss.

**Author summary:** Despite major technological and computational advances, we remain unable to match human auditory perception using machines, or to restore normal-hearing communication for those with sensorineural hearing loss. An overarching reason for these limitations is that the neural correlates of auditory perception, particularly for complex everyday sounds, remain largely unknown. Although neural responses can be measured in humans noninvasively and compared with perception, these evoked responses lack the anatomical and physiological specificity required to reveal underlying neural mechanisms. Single-unit spike-train responses can be measured from preclinical animal models with well-specified pathology; however, the disparate response types (point-process versus continuous-valued signals) have limited application of the same advanced signal-processing analyses to single-unit and evoked responses required for direct comparison. Here, we fill this gap with a unifying framework for analyzing both spike-train and evoked neural responses using advanced spectral analyses of both the slow and rapid response components that are known to be perceptually relevant for speech and music, particularly in challenging listening environments. Numerous benefits of this framework are demonstrated here, which support its potential to advance the translation of spike-train data from animal models to improve clinical diagnostics and technological development for real-world listening.

## Introduction

Normal-hearing listeners demonstrate excellent acuity while communicating in complex environments. In contrast, hearing-impaired listeners often struggle in noisy situations, even with state-of-the-art intervention strategies (e.g., digital hearing aids). In addition to improving our understanding of the auditory system, the clinical outcomes of these strategies can be improved by studying how the neural representation of complex sounds relates to perception in normal and impaired hearing. Numerous electrophysiological studies have explored the neural representation of perceptually relevant sounds in humans using evoked far-field recordings, such as frequency following responses (FFRs) and electroencephalograms (Clinard et al., 2010; Kraus et al., 2017; Tremblay et al., 2006). Note that we use *electrophysiology* and *neurophysiology* to refer to evoked far-field responses and single-unit responses, respectively (see S1 Table for glossary). While these evoked responses are attractive because of their clinical viability, they lack anatomical and physiological specificity. Moreover, the underlying sensorineural hearing loss pathophysiology is typically uncertain in humans. In contrast, laboratory-controlled animal models of various pathologies can provide specific neural correlates of perceptual deficits that humans experience, and thus hold great scientific and translational (e.g., pharmacological) potential. In order to synergize the benefits of both these approaches to advance basic-science and translational applications to real-world listening, two major limitations need to be addressed.

First, there exists a significant gap in relating spike-train data recorded invasively from animals and evoked noninvasive far-field recordings feasible in humans (and animals) because the two signals are fundamentally different in form (i.e., binary-valued point-process data versus continuous-valued signals). While the continuous nature of the evoked-response amplitude allows for any of the advanced signal-processing techniques developed for continuous-valued signals to be applied [e.g., multitaper approaches to robust spectral estimation (Thomson, 1982)], spike-train analyses have been much more limited (e.g., in their application to real-world signals, as reviewed in S1 Text). This is a critical gap because most perceptual deficits and machine-hearing limits occur for speech in noise rather than for speech in quiet (Moore, 2007; Scharenborg, 2007). For example, classic neurophysiological studies have quantified the temporal coding of stationary and periodic stimuli using metrics such as vector strength [VS (Goldberg and Brown, 1969; Joris and Yin, 1992; Rees and Palmer, 1989)], whereas more recent correlogram analyses have provided temporal-coding metrics for nonperiodic stimuli, such as noise (Joris et al., 2006; Louage et al., 2004). However, as reviewed in S1 Text these metrics can be influenced by distortions from nonlinear cochlear processes (Heinz and Swaminathan, 2009; Young and Sachs, 1979), and often ignore response phase information that is likely to be perceptually relevant for simple tasks (Colburn et al., 2003) as well as for speech intelligibility (Paliwal and Alsteris, 2003; Relanõ-Iborra et al., 2016).

A second important gap exists because current spectrotemporal tools to evaluate temporal coding in the auditory system are largely directed at processing of stationary signals by linear and time-invariant systems. However, the auditory system exhibits an array of nonlinear (e.g., two-tone suppression, compressive gain, and rectification) and time-varying (e.g., adaptation and efferent feedback) mechanisms (Heil and Peterson, 2015; Sayles and Heinz, 2017). These mechanisms interact with nonstationary stimulus features (e.g., frequency transitions and time-varying intensity fluctuations, Figs 1A and 1B) to shape the neural coding and perception of these signals (Delgutte, 1997; Hillenbrand and Nearey, 1999; Nearey and Assmann, 1986). In fact, the response of an auditory-nerve (AN) fiber to even a simple stationary tone shows nonstationary features, such as a sharp onset and adaptation (Fig 1C), illustrating the need for nonstationary analyses of temporal coding. However, the extensive single-unit speech coding studies using classic spike-train metrics have typically been limited to synthesized and stationary speech tokens, which has deferred the study of the rich kinematics present in natural speech (Delgutte, 1980; Sinex and Geisler, 1983; Young and Sachs, 1979). Some windowing-based approaches have been used to study time-varying stimuli and responses (Cariani and Delgutte, 1996a; Sayles and Winter, 2008), but the approaches used have imposed a limit on the temporal and spectral resolution with which dynamics of the auditory system can be studied.

**Fig 1.**
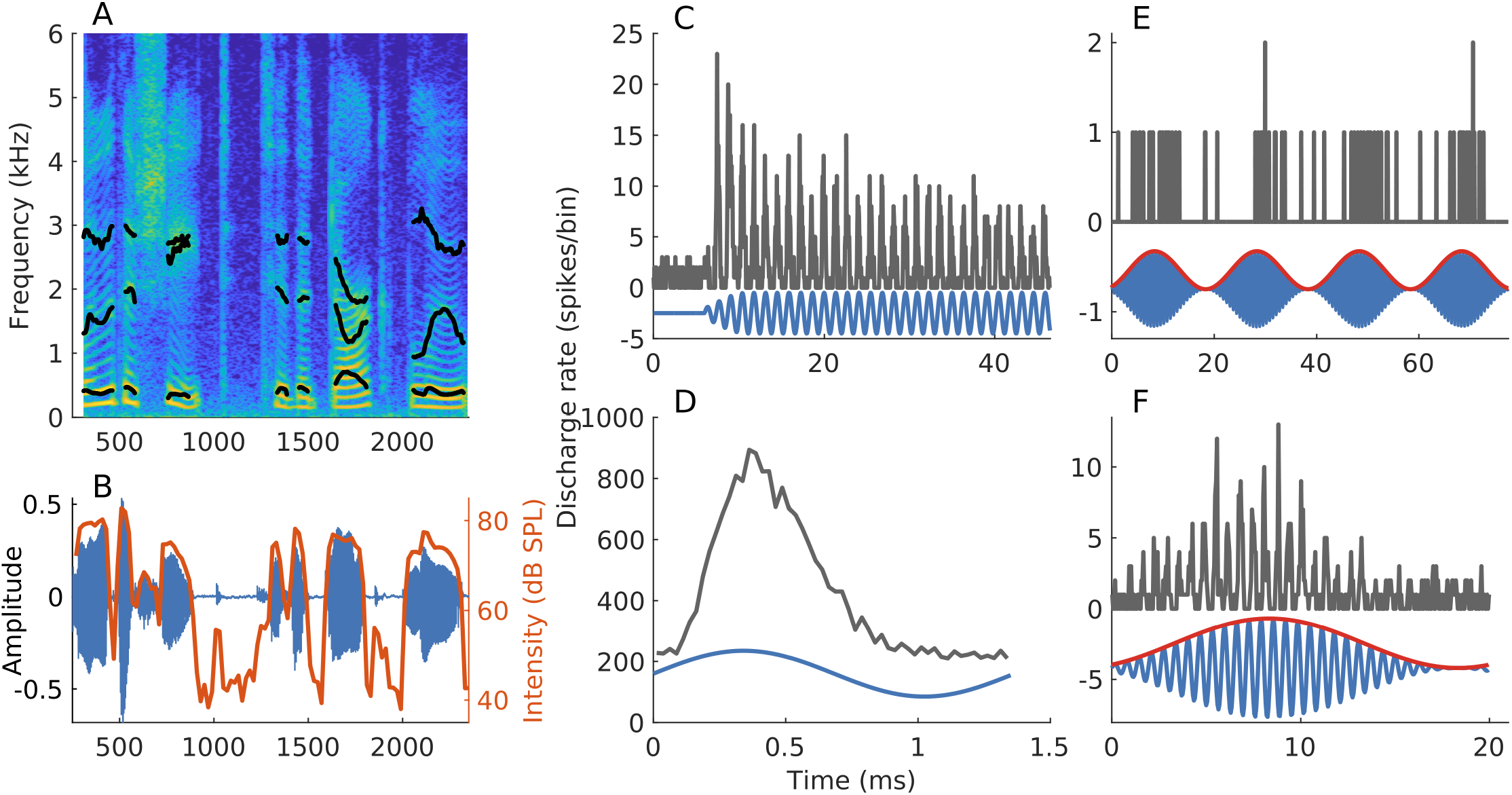
Neural responses of AN fibers are invariably nonstationary, even when the stimulus is not. (A, B) Spectrogram and waveform of a speech segment (*s*_4_ described in *Materials and Methods*). Formant trajectories (black lines in panel A) and short-term intensity (red line in panel B, computed over 20-ms windows with 80% overlap) vary with time, highlighting two nonstationary aspects of speech stimuli. (C) PSTH constructed using spike trains in response to a tone at the AN-fiber’s characteristic frequency [CF, most-sensitive frequency; fiber had CF=730 Hz, and was high spontaneous rate or SR (Liberman, 1978)]. Tone intensity = 40 dB SPL. Even though the stimulus is stationary, the response is nonstationary (i.e., sharp onset followed by adaptation). (D) Period histogram, constructed from the data used in C, demonstrates the phase-locking ability of neurons to individual stimulus cycles. (E) PSTH constructed using spike trains in response to a sinusoidally amplitude-modulated (SAM) CF-tone (50-Hz modulation frequency, 0-dB modulation depth, 35 dB SPL) from an AN fiber (CF = 1.4 kHz, medium SR). (F) Period histogram (for one modulation period) constructed from the data used in E. The response to the SAM tone follows both the modulator (envelope, red, panels E and F) as well as the carrier (temporal fine structure), the rapid fluctuations in the signal (blue, panel F). Bin width = 0.5 ms for histograms in C-F. Number of stimulus repetitions for C and E were 300 and 16, respectively.

The present study focuses on developing spectrotemporal tools to characterize the neural representation of kinematics naturally present in real-world signals, speech in particular, that are appropriate for the nonlinear and time-varying auditory system. We describe a unifying framework to study temporal coding in the auditory system, which allows direct comparison of single-unit spike-train responses with evoked far-field recordings. In particular, we demonstrate the unifying merit of using alternating-polarity peristimulus time histograms (*apPSTHs*, Table 1), a collection of PSTHs obtained from responses to both positive and negative polarities of the stimulus. By using both polarities, neural coding of natural sounds can be studied using the common temporal dichotomy between the slowly varying envelope (ENV) and rapidly varying temporal fine structure (TFS) (Figs 1E and 1F), which has been especially relevant for speech-perception studies (Shannon et al., 1995; Smith et al., 2002). We derive explicit relations between *apPSTHs* and existing metrics for quantifying temporal coding in auditory neurophysiology (reviewed in S1 Text), namely VS and correlograms, to show that no information is lost by using *apPSTHs*. In fact, the use of *apPSTHs* is computationally more efficient, provides more precise spectral estimators, and opens up new avenues for perceptually relevant analyses that are otherwise not possible. Next, an *apPSTH* -based ENV/TFS taxonomy is presented, including existing and new metrics. This taxonomy allows for spectrally specific analyses that avoid analysis distortions due to inner-hair-cell transduction and synaptic rectification processes, resulting in more accurate characterizations of temporal coding than with previous metrics. Finally, these methods are extended in novel ways to include the study of nonstationary signals at superior spectrotemporal resolution compared to conventional windowing-based approaches, like the spectrogram or wavelet analysis.

**Table 1.**
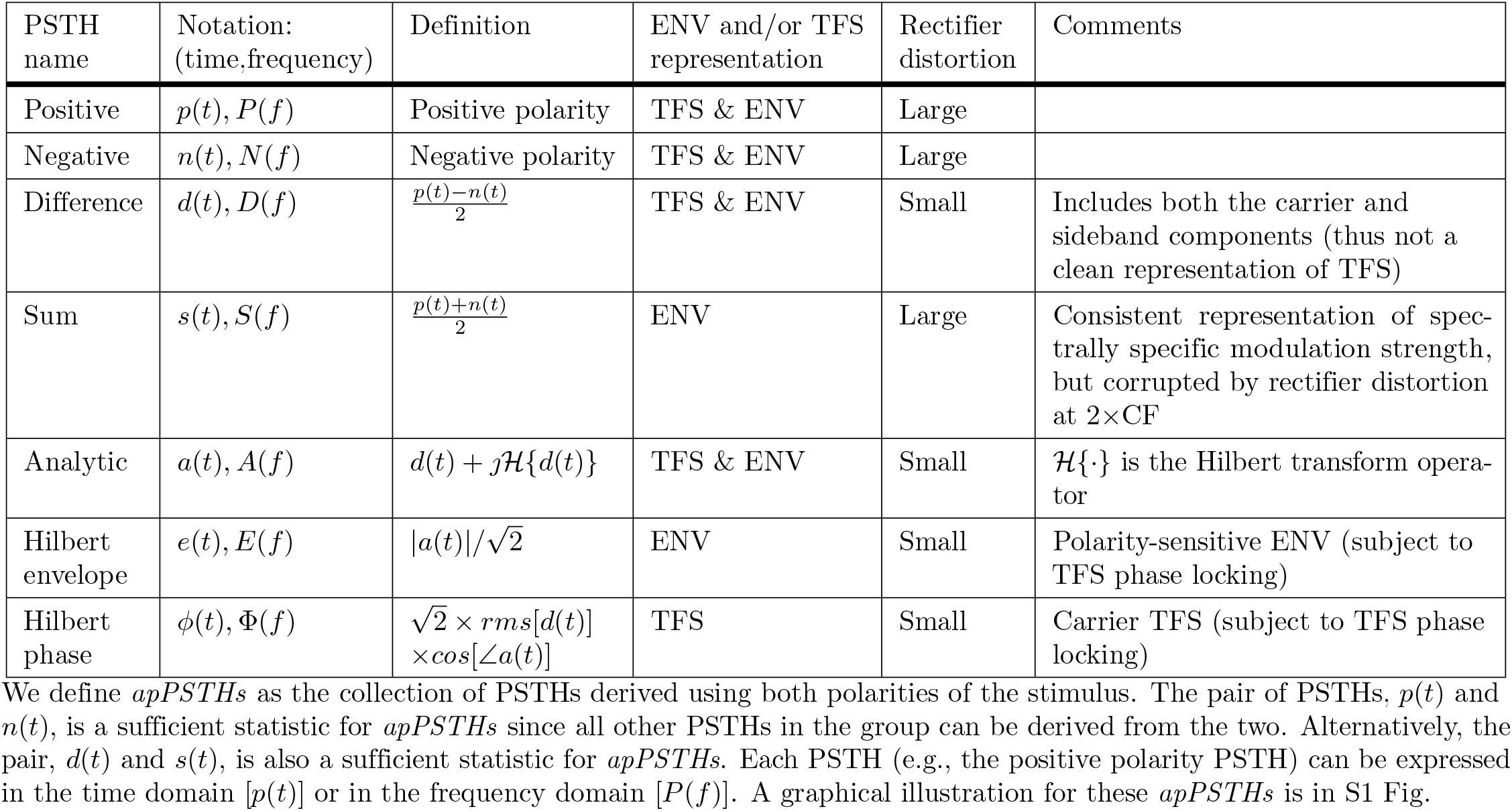
*apPSTH* -taxonomy for ENV & TFS.

### A unified framework for quantifying temporal coding based on alternating-polarity PSTHs (*apPSTHs*)

In this section, we first show that *apPSTHs* can be used to unify classic metrics, e.g., *VS* and correlograms (reviewed in S1 Text), in a computationally efficient manner. Then, we show that *apPSTHs* offer more precise spectral estimates compared to correlograms, and allow for perceptually relevant analyses that are not possible with classic metrics.

### *apPSTHs* permit computationally efficient temporal analyses

Let us denote the PSTHs in response to the positive and negative polarities of a stimulus as *p*(*t*) and *n*(*t*), respectively. Then, the *sum PSTH, s*(*t*), which represents the polarity-tolerant component in the response, is estimated as

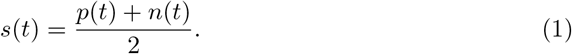

The *difference PSTH, d*(*t*), which represents the polarity-sensitive component in the response, is estimated as

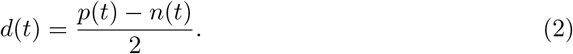

The difference PSTH has been previously described as the compound PSTH(Goblick and Pfeiffer, 1969). Here we use the terms *sum* and *difference* for *s*(*t*) and *d*(*t*), respectively, for clarity. Compared to the spectra of the single-polarity PSTHs [i.e., of *p*(*t*) or *n*(*t*)], the spectrum of the difference PSTH, *D*(*f*), is substantially less corrupted by rectifier-distortion analysis artifacts [(Sinex and Geisler, 1983), also see S1 Fig panels B and D]. This improvement occurs because even-order distortions, which strongly contribute to these artifacts, are effectively canceled out by subtracting PSTHs for opposite polarities. A second way spectral peaks absent in the stimulus can arise in the *p*(*t*)-spectrum is because of propagating combination tones of cochlear origin [e.g., distortion products, (Kemp, 1978)]. Unlike rectifier distortion, which is an artifact of analysis, combination tones are present in the cochlea and can affect perception. As the phase of these combination tones depends on stimulus polarity (Kemp, 1978), these perceptually relevant combination tones are captured in the difference PSTH. These distinct sources are discussed in more detail by Young and Sachs with respect to analyses of stationary synthesized-vowel responses from AN fibers (Young and Sachs, 1979).

The Fourier magnitude spectrum of the difference PSTH has been referred to as the synchronized rate. We show that the synchronized rate relates to *V S* by

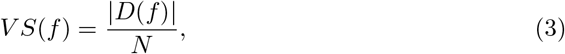

where *f* is frequency in Hz, and *N* is the total number of spikes (S2 Appendix).

In addition, we demonstrate that the autocorrelogram and the shuffled autocorrelation (SAC) function of the PSTH are related (S3 Appendix), which leads to important computational efficiencies. In particular the SAC for a set of M spike trains *X* = {*<u>x</u>*<u>_1_</u>, *<u>x</u>*<u>_2_</u>, …, *<u>x</u>*_*<u>M</u>*_} can be estimated as

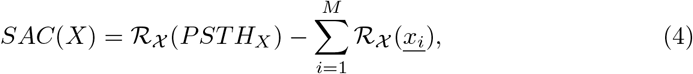

where *ℛ* _*χ*_is the autocorrelation operator, and *PSTH*_*X*_ is the PSTH constructed using *X*. Similarly, the SCC for two sets of spike trains *X* = {*<u>x</u>*<u>_1_</u>, *<u>x</u>*<u>_2_</u>, …, *<u>x</u>*_*<u>L</u>*_} and *Y* = {*<u>y</u>*<u>_1_</u>, *<u>y</u>*<u>_2_</u>, …, *<u>y</u>*_*<u>M</u>*_} can be estimated as

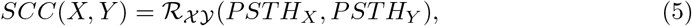

where *PSTH*_*X*_ and *PSTH*_*Y*_ are PSTHs constructed using *X* and *Y*, respectively, and ℛ_*𝒳𝒴*_ is the cross-correlation operator. Since SACs and SCCs can be computed using *apPSTHs*, it follows that *sumcor* and *difcor* can also be computed using *apPSTHs* (S4 Appendix). As *apPSTHs* can be used to compute correlograms, *apPSTHs* offer the same degree of smoothing as correlograms.

Importantly, the use of *apPSTHs* to compute correlograms is computationally more efficient compared to the existing correlogram-estimation method, i.e., by tallying all interspike intervals. For a fixed stimulus duration and PSTH resolution, estimating the autocorrelation function of the PSTH requires constant time complexity [𝒪(1)]. Thus, for *N* spikes, the SAC and SCC can be computed with 𝒪(*N*) complexity that is needed for constructing the PSTH using Eqs 4 and 5. This is substantially better than the 𝒪(*N* ^2^) complexity needed to compute the correlograms by tallying shuffled all-order interspike intervals. For example, consider a spike-train dataset that consists of 50 repetitions of a stimulus with 100 spikes per repetition. To compute the SAC using (all-order) ISIs, each spike time (5000 unique spikes) has to be compared with spike times from all other repetitions (4900 spike times). This tallying method requires 24.5×10^6^ (i.e., 5000 × 4900) operations to compute the SAC, where one operation consists of comparing two spike times and incrementing the corresponding SAC-bin by 1. In contrast, only 5000 operations are needed to construct the PSTH for 5000 (50 × 100) total spikes. The PSTH can then be used to estimate the SAC with constant time complexity. In addition to their computational efficiency, *apPSTHs* offer additional benefits for relating single-unit responses to far-field responses, for spectral estimation, and for speech-intelligibility modeling, as discussed below.

### *apPSTHs* unify single-unit and far-field analyses

The PSTH is particularly attractive because the PSTH from single neurons or a population of neurons, by virtue of being a continuous signal, can be directly compared to evoked potentials in response to the same stimulus (e.g., Fig 2). In this example, the speech sentence *s*_3_ was used to record the frequency following response (FFR) from one animal. The same stimulus was also used to record spike trains from AN fibers (N=246) from 13 animals. The mean *d*(*t*) and mean *s*(*t*) were computed by pooling PSTHs across all neurons. The difference and sum FFRs were estimated by subtracting and averaging FFRs to alternating polarities, respectively. This approach of estimating polarity-tolerant and polarity-sensitive FFR components is well established (Aiken and Picton, 2008; Ananthakrishnan et al., 2016; Shinn-Cunningham et al., 2013).

**Fig 2.**
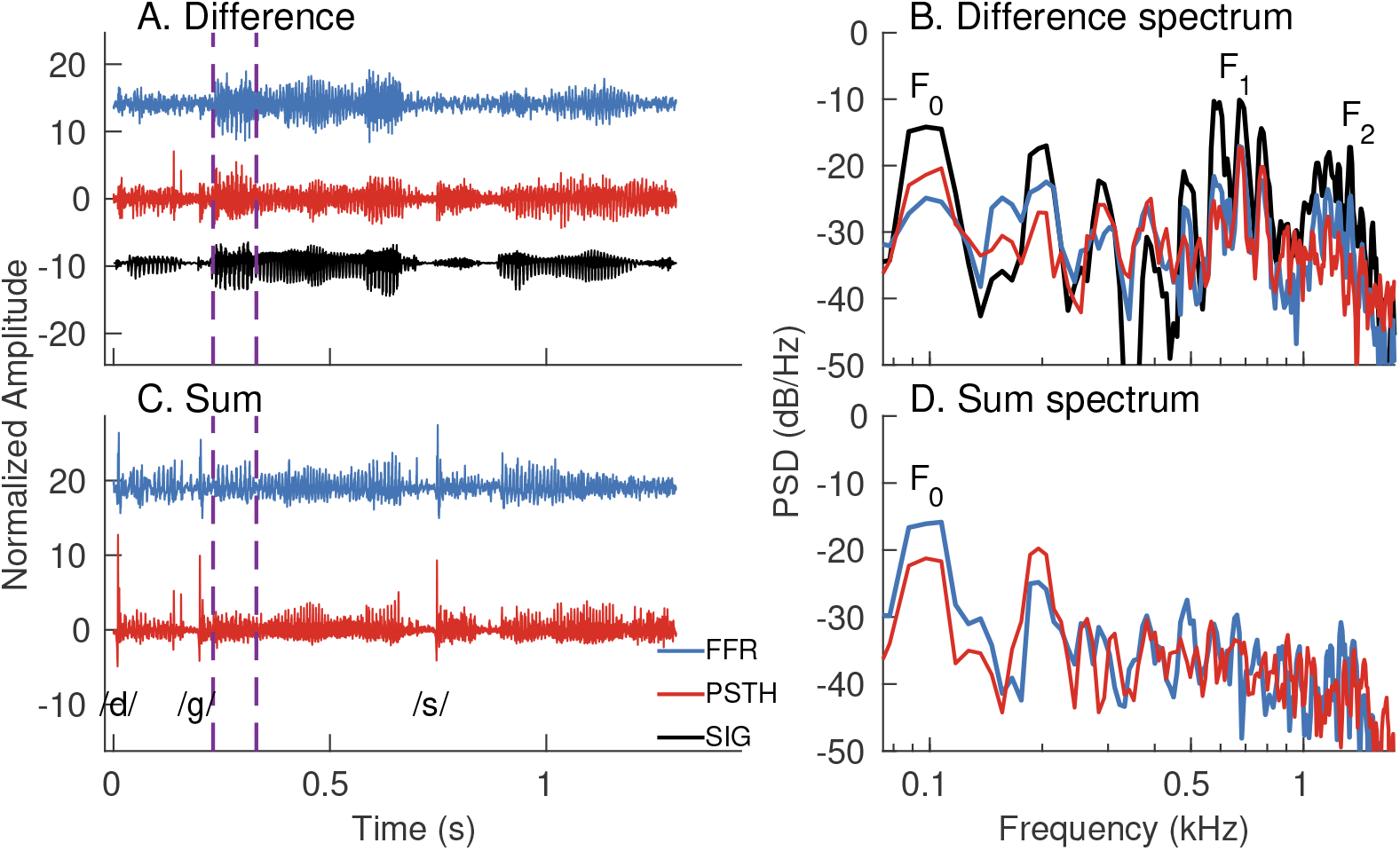
*apPSTHs* can be directly compared to evoked potentials in response to the same stimulus. (A) Time-domain waveforms for the difference FFR (blue) and mean difference PSTH [*d*(*t*), red] in response to a Danish speech stimulus, *s*_3_ (black). Mean *d*(*t*) was computed by taking the grand average of *d*(*t*)*s* from 246 AN fibers from 13 animals (CFs: 0.2 to 11 kHz). The difference FFR was estimated by subtracting FFRs to alternating stimulus polarities. (B) Spectra for the signals in A for a 100-ms segment (purple dashed lines in A). (C) Time-domain waveforms for the sum FFR (blue) and mean sum PSTH [*s*(*t*), red] for the same stimulus. Both responses show sharp onsets for plosive (/d/ and /g/) and fricative (/s/) consonants. (D) Spectra for the responses in C for the same segment considered in B. The mean *s*(*t*) was estimated as the grand average of *s*(*t*)*s* from 246 neurons. Sum FFR was estimated by halving the sum of the FFRs to both polarities. Stimulus intensity = 65 dB SPL.

Qualitatively, the periodicity information in the mean *d*(*t*) and the difference FFR were similar (Fig 2A); this is expected because the difference FFR receives significant contributions from the auditory nerve (King et al., 2016). To compare the spectra for the two responses, a 100-ms segment was considered. The first formant (*F*_1_) and the first few harmonics of the fundamental frequency (*F*_0_) were well captured in both spectra. *F*_2_ was also well captured in the difference FFR, and to a lesser extent, in the mean *d*(*t*).

The mean *s*(*t*) and the sum FFR also show comparable temporal features in these nonstationary responses (Fig 2C). For example, both responses show sharp onsets for plosive and fricative consonants. The segment considered in Fig 2B was used to compare the spectra for the two sum responses. Both spectra show similar spectral peaks near the first two harmonics of *F*_0_ (Fig 2D), which indicates that pitch-related periodicity is well captured in both the sum FFR and mean *s*(*t*). However, there are some discrepancies between the relative heights of the first two *F*_0_-harmonics. These could arise because the average FFR primarily reflects activity of high-frequency neurons from rostral generators (e.g., the inferior colliculus) (King et al., 2016), which show stronger polarity-tolerant responses compared to the auditory nerve (Joris, 2003). In contrast, the mean *s*(*t*) is based on responses of AN fibers, which show strong polarity-sensitive responses to *F*_0_ due to tuning-curve tail responses at high sound levels like that used here. These tail responses contribute to power at 2*F*_0_ as rectifier distortion. Other potential sources that can contribute to any far-field evoked response include receptor potentials (e.g., cochlear microphonic) and electrical interference. Cochlear microphonic is substantially reduced in the sum responses, although it may not be completely removed (Lichtenhan et al., 2013; Verschooten and Joris, 2014). However, cochlear microphonic should contribute to the second harmonic of the sum-FFR spectrum, and therefore does not explain the relative lack of salience for 2*F*_0_ in the sum-FFR spectrum. Electrical interference had insignificant effect on these FFR data [Fig 2 in (Parida and Heinz, 2020)]. However, in general, these sources can substantially contribute to evoked responses, such as the compound action potential, and thus should be considered when comparing these evoked responses with invasive spike-train data (Verschooten and Joris, 2014). In this regard, using the *apPSTH* –based framework to analyze invasive spike-train recordings allows direct comparison of invasive single-unit data with noninvasive continuous-valued evoked potentials and evaluation of the neural origins of evoked responses.

### Variance of *apPSTH* -based spectral estimates can be reduced relative to correlogram-based spectral estimates

Temporal information in a signal can be studied not only in the time domain (e.g., using correlograms) but also in the frequency domain (e.g., using the power spectral density, PSD). The frequency-domain representation often provides a compact alternative compared to the time-domain counterpart. In the framework of spectral estimation, the source (“true”) spectrum, which is unknown, is regarded as a parameter of a random process that is to be estimated from the available data (i.e., from examples of the random process). Spectral estimation is complicated by two factors: (1) finite response length, and (2) stochasticity of the system. The former introduces bias to the estimate, i.e., the PSD at a given frequency can differ from the true value. This bias reflects the leakage due to power at nearby (narrowband bias) and far-away (broadband bias) frequencies (due to the inherent temporal windowing from the finite-duration response). Stochasticity of the system adds randomness to the sampled data, which creates variance in the estimate. Desirable properties of PSD estimators are minimized bias and variance. Bias can be reduced by multiplying the data (prior to spectral estimation) with a taper that has a strong energy concentration near 0 Hz. Variance can be reduced by using a greater number of tapers to estimate multiple (independent) PSD estimates, which can be averaged to compute the final estimate. The multitaper approach optimally reduces the bias and variance of the PSD estimate (Babadi and Brown, 2014; Thomson, 1982). In this approach, for a given data length, a frequency resolution is chosen, based on which a set of orthogonal tapers are computed. These tapers include both even and odd tapers, which can be used to obtain the independent PSD estimates to be averaged. In contrast, for the same frequency resolution, only even tapers can be used with correlograms as they are even sequences (Oppenheim, 1999; Rangayyan, 2015). Therefore, variance in the PSD estimate can be reduced by a factor of up to 2 by using *apPSTHs* instead of correlograms.

For example, the benefit (in terms of spectral-estimation variance) of using the multitaper spectrum of *d*(*t*), as opposed to the common approach of estimating the discrete Fourier transform (DFT) of the *difcor*, can be quantified by comparing the two spectra at a single frequency (Fig 3). Here, a 100-ms segment of the *s*_3_ speech stimulus was used as the analysis window. The segment had an *F*_0_ of 98 Hz and *F*_1_ of 630 Hz (Fig 3A). Fig 3B shows example spectra estimated using spike trains recorded from a low-frequency AN fiber [CF = 900 Hz, SR = 81 spikes/s]. The multitaper spectrum was estimated using the MATLAB function *pmtm* [two tapers corresponding to a time-bandwidth product of 3, adaptive weights (Thomson, 1982)]. To compare variances in the two estimated spectra, fractional power at the 6th harmonic was considered, as this harmonic was closest to *F*_1_. This analysis was restricted to neurons (N=10) for which data was available for at least 75 repetitions per polarity and that had a CF between 0.3 and 2 kHz. For each neuron, 25 spike trains per polarity were chosen randomly 12 times to estimate fractional power at the 6th harmonic. The same set of spike trains were used to estimate distributions for both the *difcor* -spectrum and *D(f)*. The ratio of *difcor* -based fractional power variance to the *apPSTH* -based fractional power variance at 6*F*_0_ was *>*1 for all 10 neurons considered (Fig 3D), demonstrating the benefit of being able to compute a multitaper spectrum from *d*(*t*) compared to the *difcor* -spectrum in reducing variance. Overall, these results indicate that less data are required to achieve the same level of precision in a spectral metric based on the multitaper spectrum of an *apPSTH* compared to the same metric derived from the DFT of the correlogram.

**Fig 3.**
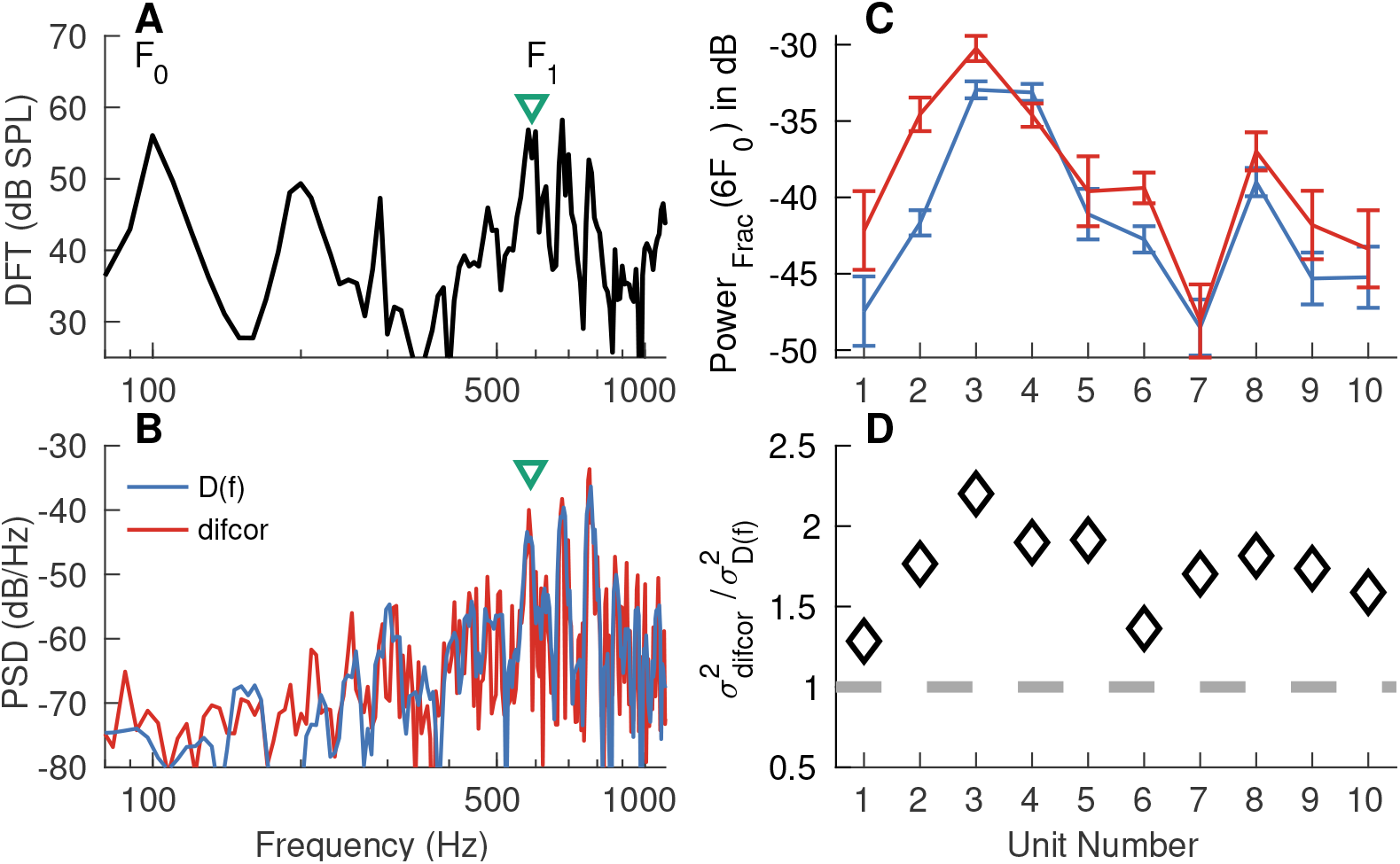
Lower spectral-estimation variance can be achieved using *apPSTHs* (with multiple tapers) compared with *difcor* correlograms. (A) Spectrum for the 100-ms segment in the speech sentence *s3* (*F*_0_ ∼ 98 Hz, *F*_1_ ∼ 630 Hz) used for analysis. (B) Example spectra for an AN fiber (CF=900 Hz, high SR) with spikes from 25 randomly chosen repetitions per polarity. The first two discrete-prolate spheroidal sequences were used as tapers corresponding to a time-bandwidth product of 3 to estimate *D*(*f*), the spectrum of *d*(*t*). No taper (i.e., rectangular window) was used to estimate the *difcor* spectrum. The AN fiber responded to the 6th, 7th and 8th harmonic of the fundamental frequency. (C) Error-bar plots for fractional power (*Power*_*Frac*_) at the frequency (green triangle) closest to the 6th harmonic. Error bars were computed for 12 randomly and independently drawn sets of 25 repetitions per polarity. The same spikes were used to compute the spectra for *d*(*t*) (blue) and *difcor* (red). (D) Diamonds denote the ratio of variances for the *difcor* -based estimate to the *d(t)*-based estimate. This ratio was greater than 1 (i.e., above the dashed gray line) for all units considered, which demonstrates that the variance for the multitaper-*d(t)* spectrum was lower than the *difcor* -spectrum variance. AN fibers with CFs between 0.3 and 2 kHz and with at least 75 repetitions per polarity of the stimulus were considered. Bin width = 0.1 ms for PSTHs. Sampling frequency = 10 kHz for FFRs. Stimulus intensity = 65 dB SPL.

### Benefits of *apPSTHs* for speech-intelligibility modeling

Speech-intelligibility (SI) models aim to predict the effects of acoustic manipulations of speech on perception. Thus, SI models allow for quantitative evaluation of the perceptually relevant features in speech. More importantly, SI models can guide the development of optimal hearing-aid strategies for hearing-impaired listeners. However, state-of-the-art SI models are largely based on the acoustic signal, where there is no physiological basis to capture the various effects of sensorineural hearing loss (SNHL) (Cooke, 2006; Houtgast and Steeneken, 1973; Kryter, 1962; Relanõ-Iborra et al., 2016; Taal et al., 2011). In contrast, neurophysiological SI models (i.e., SI models based on neural data) are particularly important in this regard since spike-train data from preclinical animal models of various forms of SNHL provide a direct way to evaluate the effects of SNHL on speech-intelligibility modeling outcomes (Heinz, 2015; Rallapalli and Heinz, 2016).

A major advantage of PSTH-based approaches over correlogram-based approaches is that they can be used to extend a wider variety of acoustic SI models to include neurophysiological data. In particular, correlograms can be used to extend power-spectrum-based SI models (Cooke, 2006; Houtgast and Steeneken, 1973; Jørgensen and Dau, 2011; Kryter, 1962; Taal et al., 2011) but not for the more recent SI models that require phase information of the response (Relanõ-Iborra et al., 2016; Scheidiger et al., 2018). For example, the speech envelope-power-spectrum model (sEPSM) has been evaluated using simulated spike trains since sEPSM only requires power in the response envelope, which can be estimated from the *sumcor* spectrum (Rallapalli and Heinz, 2016). However, *sumcor* cannot be used to evaluate envelope-phase-based SI models since it discards phase information. Studies have shown that the response phase can be important for speech intelligibility (Delgutte et al., 1998; Paliwal and Alsteris, 2003). In contrast to the *sumcor*, the time-varying PSTH contains both phase and magnitude information, and thus, can be used to evaluate both power-spectrum- and phase-spectrum-based SI models. For example, because the PSTH *p*(*t*) [or *n*(*t*)] is already rectified, it can be filtered through a modulation filter bank to estimate “internal representations” in the modulation domain (Fig 4). These spike-train-derived “internal representations” are analogous to those used in phase-spectrum-based SI models (Relanõ-Iborra et al., 2016; Scheidiger et al., 2018) and can be further processed by existing SI back-ends to estimate SI values. This example demonstrates a proof of concept of using spike-train data to evaluate a spectrally specific envelope based SI model using *apPSTHs*. In general, SI models that include a peripheral or modulation filter bank representation, which is the case for most successful SI models [e.g., the speech transmission index (Steeneken and Houtgast, 1980), the spectrotemporal modulation index (Elhilali et al., 2003), speech envelope power spectrum models (Jørgensen and Dau, 2011; Jørgensen et al., 2013)], can be evaluated using spike-train data recorded from peripheral (e.g., auditory-nerve fibers) or central (e.g., inferior colliculus) neurons, respectively, using *apPSTHs*. Therefore, these analyses allow for the evaluation of a wider variety of acoustic-based SI models in the neural domain (magnitude and phase), where translationally relevant data can be obtained from preclinical animal models of various forms of SNHL.

**Fig 4.**
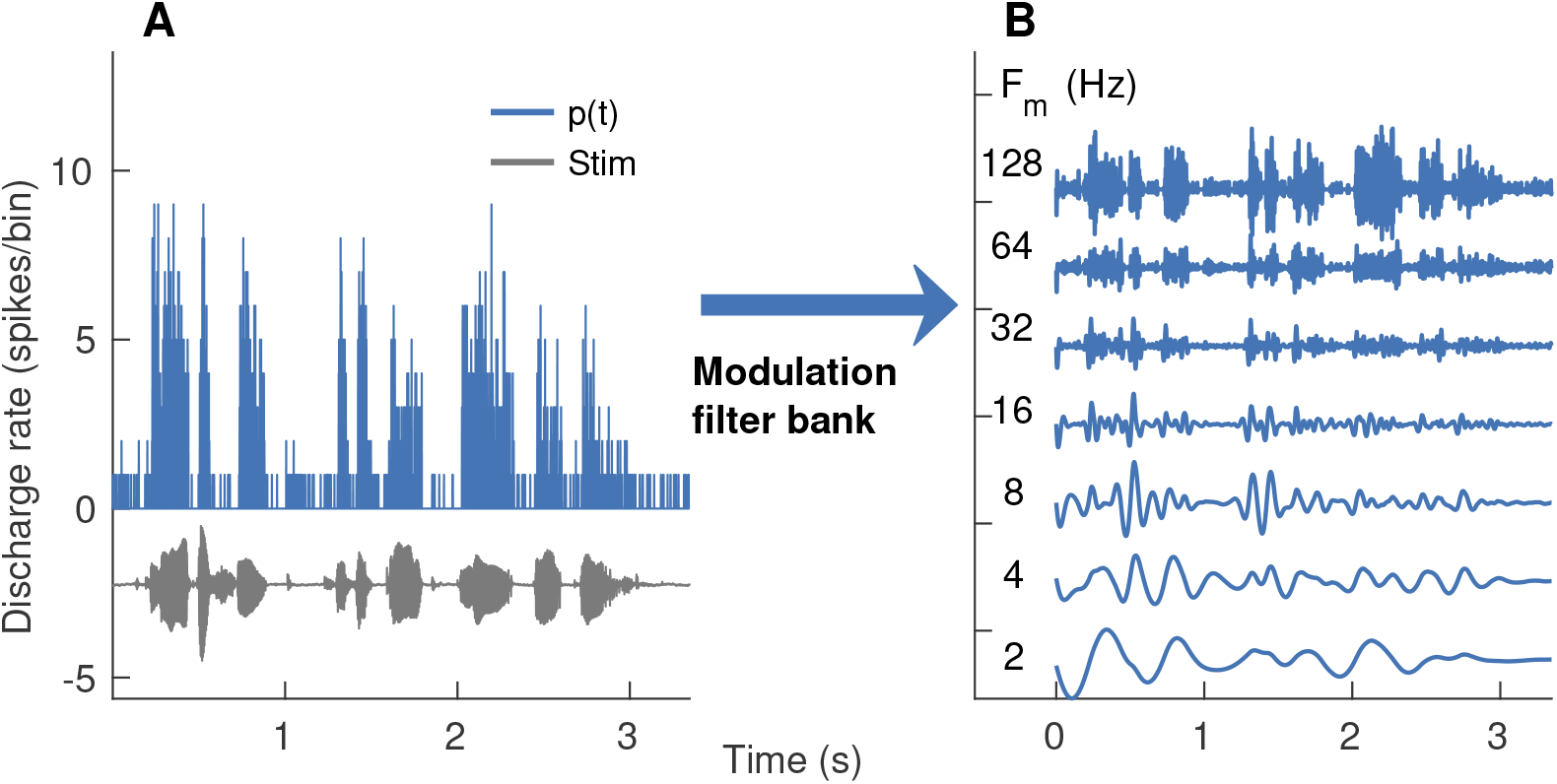
Modulation-domain internal representations for speech coding can be obtained from PSTH-based envelopes. PSTH response [*p*(*t*)] from one AN fiber (CF=290 Hz, SR= 12 spikes/s) is shown. (A) Time-domain waveforms for the stimulus (gray) and *p*(*t*) (blue). (B) Output of a modulation filter bank after the processing of *p*(*t*). Modulation filters were zero-phase, fourth-order, and octave-wide IIR filters. Center frequencies (*F*_*m*_) for these filters ranged from 2 to 128 Hz (octave spacing), similar to those used in recent psychophysically based SI models [e.g., (Relanõ-Iborra et al., 2016)]. PSTH bin width = 0.5 ms. 15 stimulus repetitions. Stimulus intensity = 60 dB SPL.

### Quantifying ENV and TFS using *apPSTHs* for stationary signals

In this section, we first describe existing and novel ENV and TFS components that can be derived from *apPSTHs*. Next, we compare relative merits of the novel components over existing ENV and TFS components using simulated data. Finally, we apply *apPSTHs* to analyze spike-train data recorded to speech and speech-like stimuli.

#### Several ENV and TFS components can be derived from *apPSTHs* with spectral specificity

The neural response envelope can be obtained from *apPSTHs* in two orthogonal ways: (1) the low-frequency signal, *s*(*t*), and (2) the Hilbert envelope of the high-frequency carrier-related energy in *d*(*t*). *s*(*t*) is thought to represent the polarity-tolerant response component, which has been defined as the envelope response (Joris, 2003; Louage et al., 2004). For a stimulus with harmonic spectrum, *s*(*t*) captures the envelope related to the beating between harmonics. In addition, onset and offset responses (e.g., in response to high-frequency fricatives, Fig 2C) are also well captured in *s*(*t*). Although *sumcor* and *s*(*t*) are related, dynamic features like onset and offset responses are captured in *s*(*t*), but not in the *sumcor* since the *sumcor* discards phase information by essentially averaging ENV coding across the whole stimulus duration. The use of sum envelope is popular in far-field responses (Aiken and Picton, 2008; Ananthakrishnan et al., 2016; Shinn-Cunningham et al., 2013) but not directly in auditory neurophysiology studies. A major disadvantage of *s*(*t*) is that it is affected by rectifier distortions if a neuron phase locks to low-frequency energy in the stimulus (e.g., Fig 5A; discussed further below).

**Fig 5.**
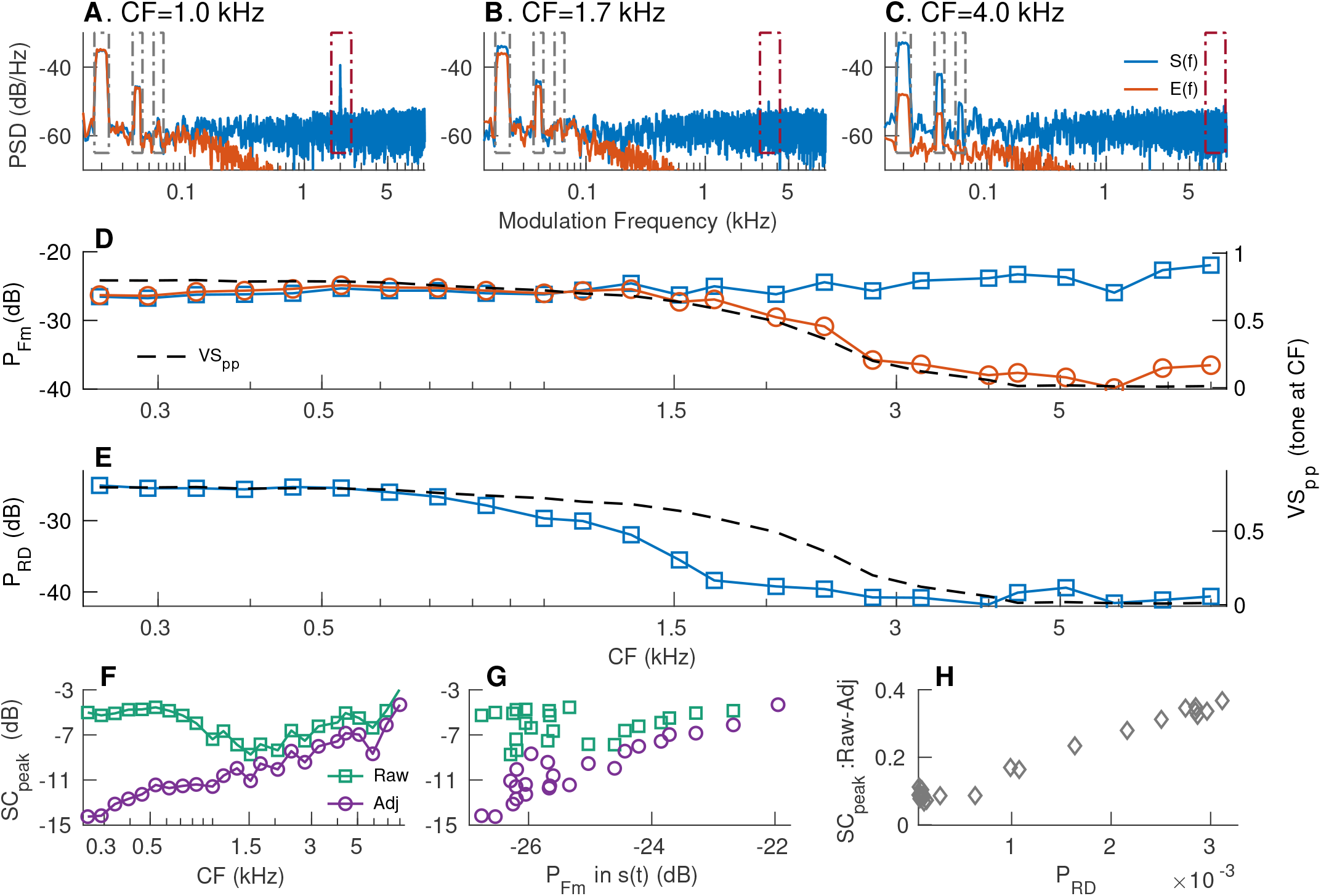
Envelope-coding metrics should be spectrally specific to avoid artifacts due to rectifier distortion and neural stochasticity. Simulated responses for 24 AN fibers (log-spaced between 250 Hz and 8 kHz) were obtained using a computational model (parameters listed in S2 Table) using SAM tones at CF (modulation frequency, *F*_*m*_=20 Hz; 0-dB (100%) modulation depth) as stimuli. Stimulus intensity ∼65 dB SPL. *S*(*f*) (blue) and *E*(*f*) (red) for three example model fibers with CFs = 1.0, 1.7, and 4 kHz (panels A-C) illustrate the relative merits of *s*(*t*) and *e*(*t*), and the potential for rectifier distortion to corrupt envelope coding metrics. *d*(*t*) was band-limited to a 200-Hz band near *F*_*c*_ for each fiber prior to estimating *e*(*t*) from the Hilbert transform of *d*(*t*). (A) For the 1-kHz fiber, *S*(*f*) and *E*(*f*) are nearly identical in the *F*_*m*_ band. *S*(*f*) is substantially affected by rectifier distortion at 2*×*CF, which can be ignored using spectrally specific analyses. (B) The two envelope spectra are largely similar near the *F*_*m*_ bands since phase-locking near the carrier (1.7 kHz) is still strong (panel D). Rectifier distortion in *S*(*f*) is greatly reduced since phase-locking at twice the carrier frequency (3.4 kHz) is weak. (C) *F*_*m*_-related power in *E*(*f*) and rectifier distortion in *S*(*f*) are greatly reduced as the frequencies for the carrier and twice the carrier are both above the phase-locking roll-off. (D) The strength of modulation coding was evaluated as the sum of the power near the first three harmonics of *F*_*m*_ (gray boxes in panels A-C) for *S*(*f*) (blue squares) and *E*(*f*) (red circles). *V S*_*pp*_ was also quantified to CF-tones for each fiber (black dashed line, right Y axis). (E) Rectifier distortion (RD) analysis was limited to the second harmonic of the carrier (brown boxes in panels A-C). RD was quantified as the sum of power in 10-Hz bands around twice the carrier frequency (2 *CF*) and the adjacent sidebands (2 × *CF* ±*F*_*m*_). RD for *E*(*f*) is not shown because *E*(*f*) was virtually free from RD. (F) Raw and adjusted *sumcor* peak-heights across CFs. *sumcors* were adjusted by band-pass filtering them in the three *F*_*m*_-related bands. Large differences between the two metrics at low frequencies indicate that the raw *sumcor* peak-heights are corrupted by rectifier distortion at these frequencies. (G) Relation between raw and adjusted *sumcor* peak-heights with *F*_*m*_-related power (from panel D) in *S*(*f*). Good correspondence between *F*_*m*_-related power in *S*(*f*) and adjusted *sumcor* peak-height supports the use of spectrally specific envelope analyses. (H) The difference between raw and adjusted *sumcor* peak-heights was largely accounted for by RD power. However, this difference was always greater than zero, suggesting broadband metrics can also be biased because of noise related to neural stochasticity.

A second way envelope information in the neural response can be quantified is by computing the envelope of the difference PSTH, *d*(*t*). This envelope, *e*(*t*), can be estimated as the magnitude of the analytic signal, *a*(*t*), of the difference PSTH

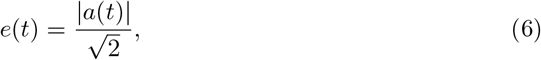

where *a*(*t*) = *d*(*t*) + _j_*ℋ* {*d*(*t*)}, and ℋ{·} is the Hilbert transform operator. The factor 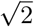 normalizes for the power difference after applying the Hilbert transform. *d*(*t*) is substantially less affected by rectifier distortion (Sinex and Geisler, 1983), and thus, so is *e*(*t*). The use of *e*(*t*) parallels the procedure followed by many computational models that extract envelopes from the output of cochlear filterbanks (Dubbelboer and Houtgast, 2008; Jørgensen and Dau, 2011; Sadjadi and Hansen, 2011).

The TFS component can also be estimated in two ways: (1) *d*(*t*), and (2) cosine of the Hilbert phase of *d*(*t*). The difference PSTH has been traditionally called the TFS response because it is the polarity-sensitive component. *difcor* and derived metrics relate to *d*(*t*) as the *difcor* is related to the autocorrelation function of *d*(*t*) (S4 Appendix). However, *d*(*t*) does not represent the response to only the carrier (phase) since it also contains envelope information in *e*(*t*). We propose a novel representation of the TFS response component, *ϕ*(*t*), estimated as the cosine phase of the analytic signal

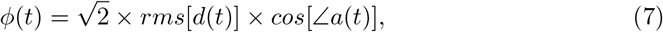

where normalization by 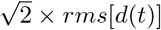 is used to match the power in *ϕ*(*t*) with the power in *d*(*t*) since *cos*[∠*a*(*t*)] is a constant-rms 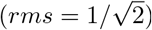 signal.

### Relative merits of sum and Hilbert-envelope PSTHs in representing spike-train envelope responses

The relative merits of the two envelope PSTHs, *s*(*t*) and *e*(*t*), were evaluated based on simulated spike-train data generated using a computational model of AN responses (Bruce et al., 2018). The model includes both cochlear-tuning and hair-cell transduction nonlinearities in the auditory system. Modulation spectra for sinusoidal amplitude-modulated (SAM) tones were estimated for *s*(*t*) and *e*(*t*) [denoted by *S*(*f*) and *E*(*f*), respectively] for individual-fiber responses (Figs 5A-5C). *d*(*t*) was band-pass filtered near CF (200-Hz bandwidth, 2nd order filter) before applying the Hilbert transform to minimize the spectral energy in *d*(*t*) that was not stimulus related. The two envelopes were evaluated based on their representations of the modulator and rectifier distortion. Rectifier distortions are expected to occur at even multiples of the carrier and nearby sidebands (i.e., 2*nF*_*c*_, 2*nF*_*c*_ − *F*_*m*_, and 2*nF*_*c*_ + *F*_*m*_ for integers n, Fig 5A). It is desirable for an envelope metric to consistently represent envelope coding across CFs and be less affected by rectifier-distortion artifacts. Modulation coding for the simulated responses was quantified as the power in 10-Hz bands centered at the first three harmonics of *F*_*m*_ (i.e., 15 to 25 Hz, 35 to 45 Hz, and 55 to 65 Hz) for both *s*(*t*) and *e*(*t*) (Fig 5D). The need to include multiple harmonics of *F*_*m*_ arises because the response during a stimulus cycle departs from sinusoidal shape due to the saturating nonlinearity associated with inner-hair-cell transduction (S2 Fig). While *F*_*m*_-related power was nearly constant across CF for *s*(*t*), it was nearly constant for *e*(*t*) only up to 1.2 kHz, after which it rolled off. This roll-off for *e*(*t*) is not surprising since *e*(*t*) relies on phase-locking near the carrier and the sidebands, as confirmed by the strong correspondence between tonal phase-locking at the carrier frequency and *F*_*m*_-related power in *e*(*t*) (Fig 5D).

The analysis of rectifier distortion was limited to only the distortion components near the second harmonic of the carrier (i.e., 2*F*_*c*_, 2*F*_*c*_ − *F*_*m*_, and 2*F*_*c*_ + *F*_*m*_) since this harmonic is substantially stronger than higher harmonics (e.g., Fig 5A). Rectifier distortion was quantified as the sum of power in 10-Hz bands centered at the three distortion frequency components. Because *e*(*t*) was estimated from spectrally specific *d*(*t*), which was band-limited to 200 Hz near the carrier frequency, *e*(*t*) was virtually free from rectifier distortion. In contrast, *s*(*t*) was substantially affected by rectifier distortion for simulated fibers with CFs below ∼2 kHz (Fig 5E). Rectifier distortion in *S*(*f*) dropped for fibers with CF above ∼0.8 kHz because phase locking at distortion frequencies (i.e., twice the carrier frequencies) was attenuated by the roll-off in tonal phase locking. For example, the simulated AN fiber in Fig 5B (CF = 1.7 kHz) maintained comparable *F*_*m*_-related power for both envelopes, but rectifier distortion for *s*(*t*) was substantially diminished because the distortion frequency (3.4 kHz) is well above the phase-locking roll-off. These results indicate that *s*(*t*) is substantially corrupted by rectifier distortion (at twice the stimulus frequency) when the neuron responds to stimulus energy that is below half the phase-locking cutoff.

Next, these spectral power metrics were compared with the correlogram-based metric, *sumcor* peak-height (Figs 5F-5H). The *sumcor* peak-height metric is defined as the maximum value of the normalized time-domain *sumcor* function (Louage et al., 2004). Prior to estimating the peak-height, the *sumcor* is sometimes adjusted by adding an inverted triangular window to compensate for its triangular shape (Heinz and Swaminathan, 2009). Here, *sumcors* were compensated by subtracting a triangular window from it so that the baseline *sumcor* is a flat function with a value of 0 (instead of 1) in the absence of ENV coding. In S5 Appendix, we show that the *sumcor* peak-height is a broadband metric and it is related to the total power in *s*(*t*), including rectifier distortions. When the *sumcor* is used to analyze responses of low-frequency AN fibers to broadband noise stimuli, the *sumcor* -spectrum, and thus, the *sumcor* peak-height, are corrupted by rectifier distortions (Heinz and Swaminathan, 2009). Similar to *S*(*f*) for low-frequency SAM tones (Fig 5A), these distortions show up at 2 ×CF in the *sumcor* -spectrum, whereas the *difcor* -spectrum has energy only near CF (Heinz and Swaminathan, 2009). Heinz and colleagues addressed these distortions by low-pass filtering the *sumcor* below CF to remove the effects of rectifier distortion at 2× CF. Here, we generalize this issue by comparing the *sumcor* and spectrally specific ENV metrics for narrowband SAM-tone stimuli to demonstrate the limitations of any broadband ENV metric. *sumcors* were adjusted by band-limiting them to 10-Hz bands near the first three harmonics of *F*_*m*_. As expected, the difference between the raw and adjusted *sumcor* peak-heights was large at low CFs (Fig 5F), where rectifier distortion corrupts the broadband *sumcor* peak-height metric. At high CFs (above 1.5 kHz), the difference between raw and adjusted *sumcor* peak-heights was small but nonzero. These differences correspond to power in *S*(*f*) at frequencies other than the modulation-related bands and reflect the artifacts of neural stochasticity due to finite number of stimulus trials. As power is always nonnegative, including power at frequencies unrelated to the target frequencies adds bias and variance to any broadband metric. The adjusted *sumcor* peak-height, unlike the raw *sumcor* peak-height, showed good agreement with spectrally specific *F*_*m*_-related power in *S*(*f*) (Fig 5G).

Overall, these results support the use of spectrally specific analyses to quantify ENV coding in order to minimize artifacts due to rectifier distortion as well as the effects of neural stochasticity. Of the two candidate *apPSTHs* to quantify response envelope, *e*(*t*) had the benefit of minimizing rectifier distortion. However, *e*(*t*)’s reliance on carrier-related phase locking limits the use of *e*(*t*) as a unifying ENV metric across the whole range of CFs. Instead, spectrally specific *s*(*t*) is more attractive because of its robustness in representing the response envelope across CFs (Fig 5D).

### Relative merits of difference and Hilbert-phase PSTHs in representing spike-train TFS responses

In order to evaluate the relative merits of *d*(*t*) and *ϕ*(*t*) in representing the neural TFS response, the same set of simulated AN spike-train responses were used as in Fig 5. Although the stimulus has power at the carrier (*F*_*c*_) and sidebands (*F*_*c*_ ± *F*_*m*_; 6 dB lower), only the carrier representation should be considered towards quantifying the TFS response because the energy at the sidebands arises due to the modulation of the carrier by the modulator (ENV). As the carrier has energy at a single frequency (*F*_*c*_) for a SAM tone, the desirable TFS response should have maximum energy concentrated at the carrier frequency and not the sidebands. Therefore, the merits of *d*(*t*) and *ϕ*(*t*) were evaluated based on how well they capture the carrier and suppress the sidebands (Fig 6).

**Fig 6.**
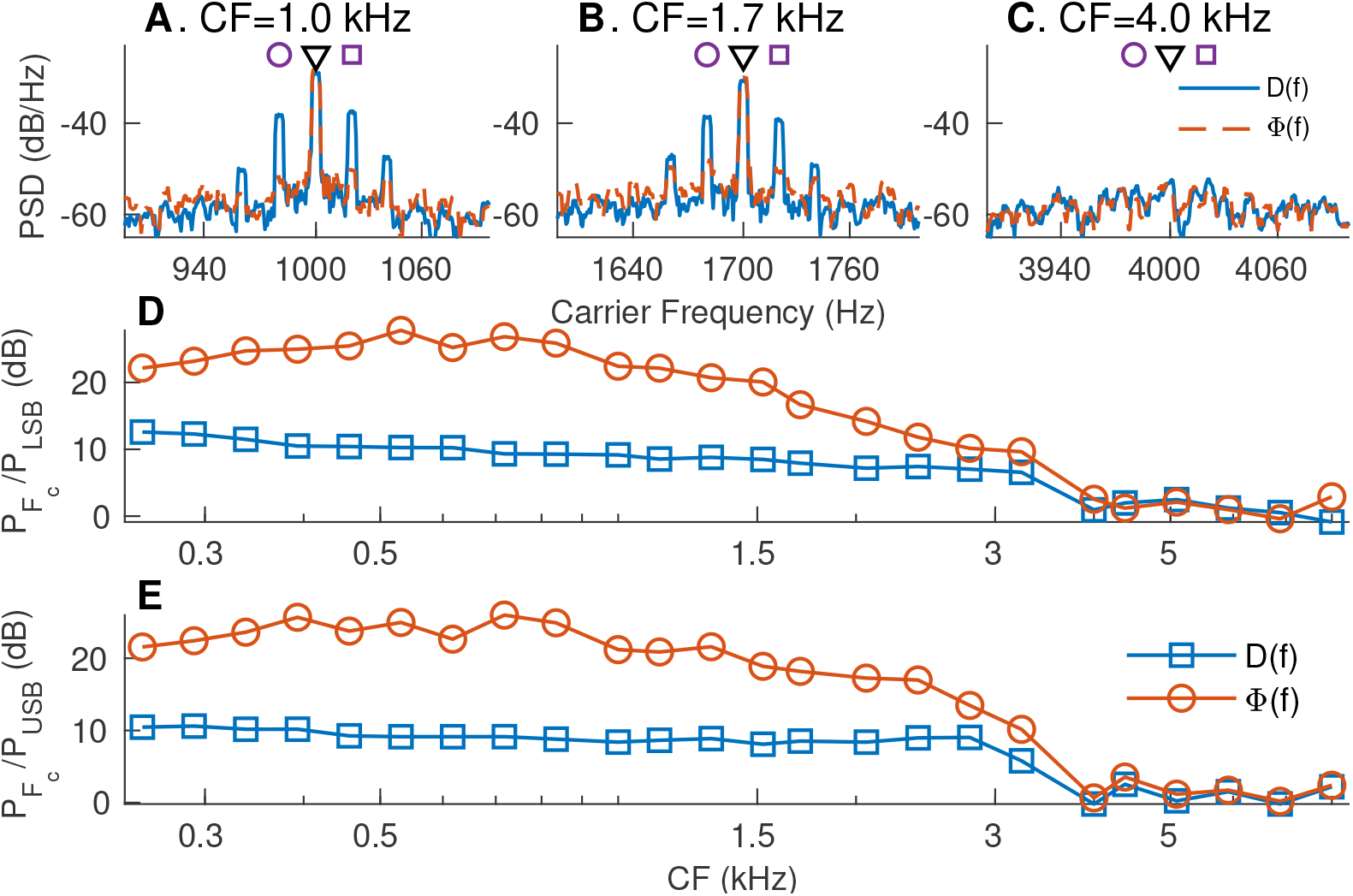
Compared to the *d*(*t*), the *apPSTH ϕ*(*t*) provides a better TFS representation. (A-C) Spectra of *d*(*t*) and *ϕ*(*t*) for the same three simulated AN fiber responses for which ENV spectra were shown in Fig 5. *D*(*f*) has substantial power at CF (black triangle), as well as at lower (purple circle) and upper (purple square) sidebands. Φ(*f*), the spectrum of *ϕ*(*t*), shows maximum power concentration at *CF* (carrier frequency), with greatly reduced sidebands. (D) Ratio of power at CF (carrier, black triangle in panels A-C) to power at lower sideband (LSB, *F*_*c*_ − *F*_*m*_, purple circles in panels A-C). (E) Ratio of power at CF (carrier) to power at upper sideband (USB, *F*_*c*_ + *F*_*m*_, purple squares in A-C). *ϕ*(*t*) highlights the carrier and not the sidebands, and thus, compared to *d*(*t*), *ϕ*(*t*) is a better representation of the true TFS response.

As mentioned previously, *d*(*t*) was band-limited to a 200-Hz bandwidth near the carrier frequency before estimating *ϕ*(*t*). *D*(*f*) at low CFs contained substantial energy at both the carrier and the sidebands (Figs 6A and 6B). This indicates that *d*(*t*) represents the complete neural coding of the SAM tone (both the envelope and the carrier) and not just the carrier. Furthermore, *D*(*f*) has additional sidebands (*F*_*c*_ ± 2*F*_*m*_) around the carrier frequency. These sidebands arise as a result of the saturating nonlinearity associated with inner-hair-cell transduction (S2 Fig), and thus, should not be considered towards TFS response. In contrast, Φ(*f*), the spectrum of *ϕ*(*t*) had most of its power concentrated at the carrier frequency, with substantially less power in the sidebands (Figs 6A and 6B). These results were consistent across a wide range of CFs and for both sidebands (Figs 6D and 6E). Overall, these results show that *ϕ*(*t*) is a better PSTH compared to *d*(*t*) in quantifying the response TFS since *ϕ*(*t*) emphasizes power at the carrier frequency and not at the sidebands.

In the following, we apply *apPSTH* -based analyses on spike-train data recorded from chinchilla AN fibers in response to speech and speech-like stimuli. In these examples, we particularly focus on certain ENV features, such as pitch coding for vowels and response onset for consonants, and TFS features, such as formant coding for vowels.

### Neural characterization of ENV and TFS using *apPSTHs* for a natural speech segment

Most previous studies have used the period histogram to study speech coding in the spectral domain (Delgutte and Kiang, 1984a; Young and Sachs, 1979). The period histogram is limited to stationary periodic stimuli, which were employed in those studies. In contrast, the use of *apPSTHs* facilitates the spectral analysis of neural responses to natural speech stimuli, which need not be stationary. Fig 7 shows the response spectra obtained using various *apPSTHs* [*p*(*t*), *s*(*t*), *d*(*t*), and *ϕ*(*t*)] for a low-frequency AN fiber in response to a natural speech segment [see S3 Fig for similar analyses for synthesized speech demonstrating the well-known “synchrony-capture” phenomenon (Delgutte and Kiang, 1984a; Young and Sachs, 1979)]. In this example, the response of a low-frequency AN fiber to a 100-ms vowel segment of the *s*_3_ natural speech sentence was considered. The CF (1.1 kHz) of this neuron is close to the second formant (*F*_2_) of this segment (Fig 7B). *P* (*f*) shows peaks corresponding to *F*_2_ (∼1.2 kHz) and *F*_0_ (∼130 Hz, Fig 7C). Similar to S3 Fig, both *D*(*f*) and Φ(*f*) show substantial energy near the formant closest to the neuron’s CF. In contrast to S3 Fig, *S*(*f*) [and *E*(*f*)] shows substantial energy near the fundamental frequency (inconsistent with synchrony capture). A detailed discussion of this discrepancy is beyond the scope of the present report, except to say that this lack of synchrony capture for natural vowels is a consistent finding that will be reported in a future study. The presence of substantial energy near *F*_0_ in *E*(*f*) indicates that *d*(*t*) is corrupted by pitch-related modulation in *e*(*t*). This is because, mathematically, *D*(*f*) is the convolution of the true TFS spectrum [Φ(*f*)] and the Hilbert-envelope spectrum [*E*(*f*)]. Overall, these results demonstrate the application of various *apPSTHs* to study the neural representation of natural nonstationary speech stimuli in the spectral domain.

**Fig 7.**
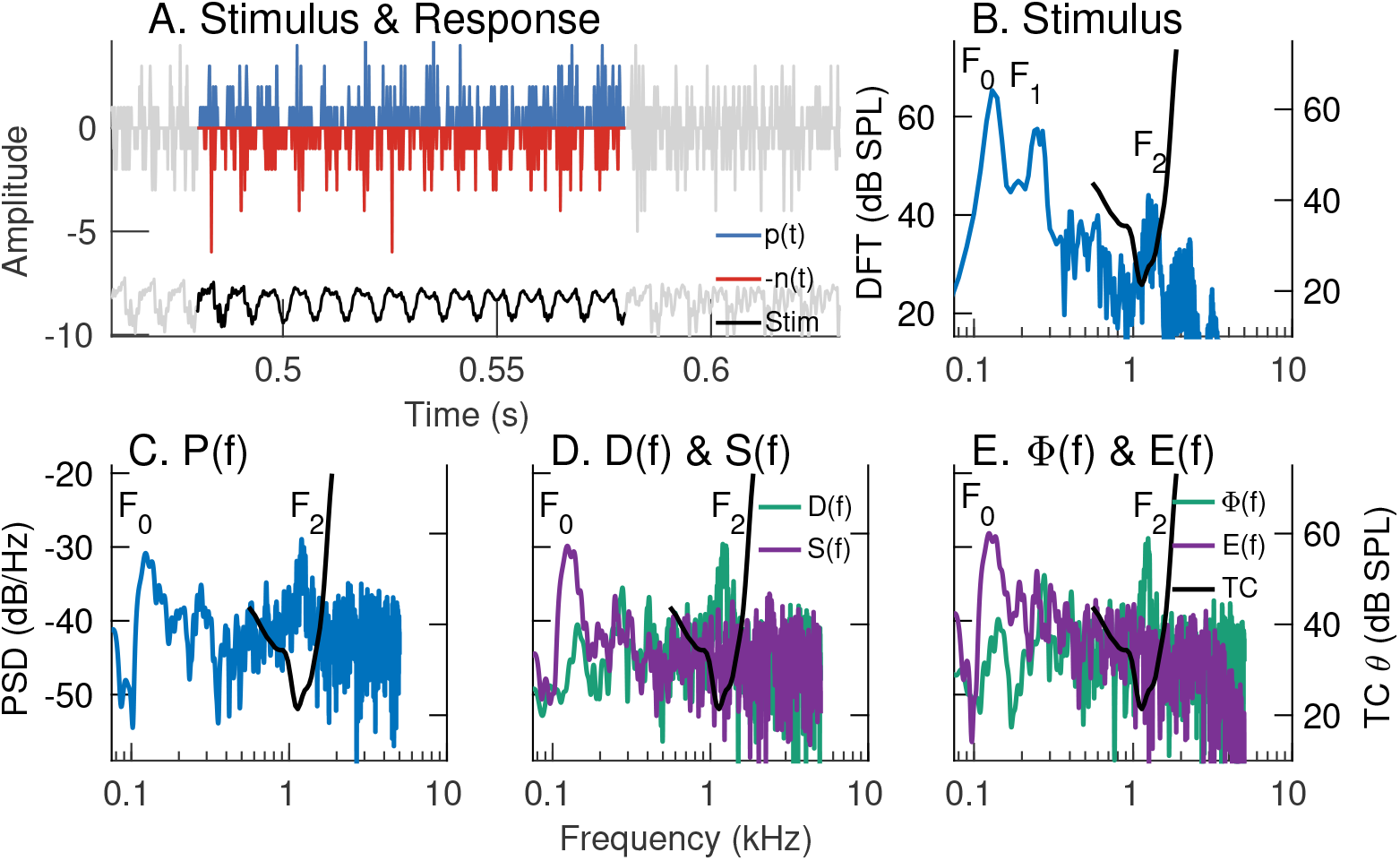
Spectral-domain application of various *apPSTHs* to spike trains recorded in response to natural speech. Example of spectral analyses of spike trains recorded from an AN fiber (CF= 1.1 kHz, SR=64 spikes/s) in response to a vowel snippet of a speech stimulus (*s*_3_). (A) Time-domain representation of *p*(*t*), *n*(*t*), and the stimulus (*Stim*). *n*(*t*) is flipped along the y-axis for display. Signals outside the analysis window are shown in gray. PSTH bin width = 0.1 ms. Number of stimulus repetitions per polarity = 50. Stimulus intensity = 65 dB SPL. (B) Stimulus spectrum (blue, left yaxis). In panels B-E, the frequency-threshold tuning curve (TC *θ*, black) of the neuron is plotted on the right y-axis. (C) *P* (*f*), which shows comparable energy at *F*_0_ (130 Hz) and *F*_2_ (1.2 kHz). (D) *D*(*f*) and *S*(*f*). (E) Φ(*f*) and *E*(*f*). Both *S*(*f*) and *E*(*f*) show peaks near *F*_0_. Similarly, both *D*(*f*) and Φ(*f*) show good *F*_2_ representations, although *D*(*f*) is corrupted by the strong *F*_0_-related modulation in *e*(*t*) as *d*(*t*) = *e*(*t*) *× ϕ*(*t*). The significant representation of *F*_0_ in this near-*F*_2_ AN fiber response to a natural vowel is inconsistent with the synchrony-capture phenomenon for synthetic stationary vowels.

### Onset envelope is well represented in the sum PSTH but not in the Hilbert-envelope PSTH

In addition to analyzing spectral features, *apPSTHs* can also be used to analyze temporal features in the neural response. An example temporal feature is the onset envelope, which has been shown to be important for neural coding of consonants (Delgutte, 1980; Heil, 2003), in particular fricatives (Delgutte and Kiang, 1984b). A diminished onset envelope in the peripheral representation of consonants is hypothesized to be a contributing factor for perceptual deficits experienced by hearing-impaired listeners (Allen and Li, 2009), and thus is important to quantify. Fig 8 shows example onset responses for a high-frequency AN fiber (CF= 5.8 kHz, SR= 70 spikes/s) for a fricative (*/s/*) portion of the speech stimulus *s*_3_. The onset is well captured in single-polarity PSTHs [*p*(*t*) and *n*(*t*), Fig 8A] and in the sum envelope [*s*(*t*), Fig 8B]. Since the onset is a polarity-tolerant feature, it is greatly reduced by subtracting the PSTHs to opposite polarities. As a result, response onset is poorly captured in *d*(*t*) (Fig 8C) and its Hilbert envelope, *e*(*t*) (Fig 8D).

**Fig 8.**
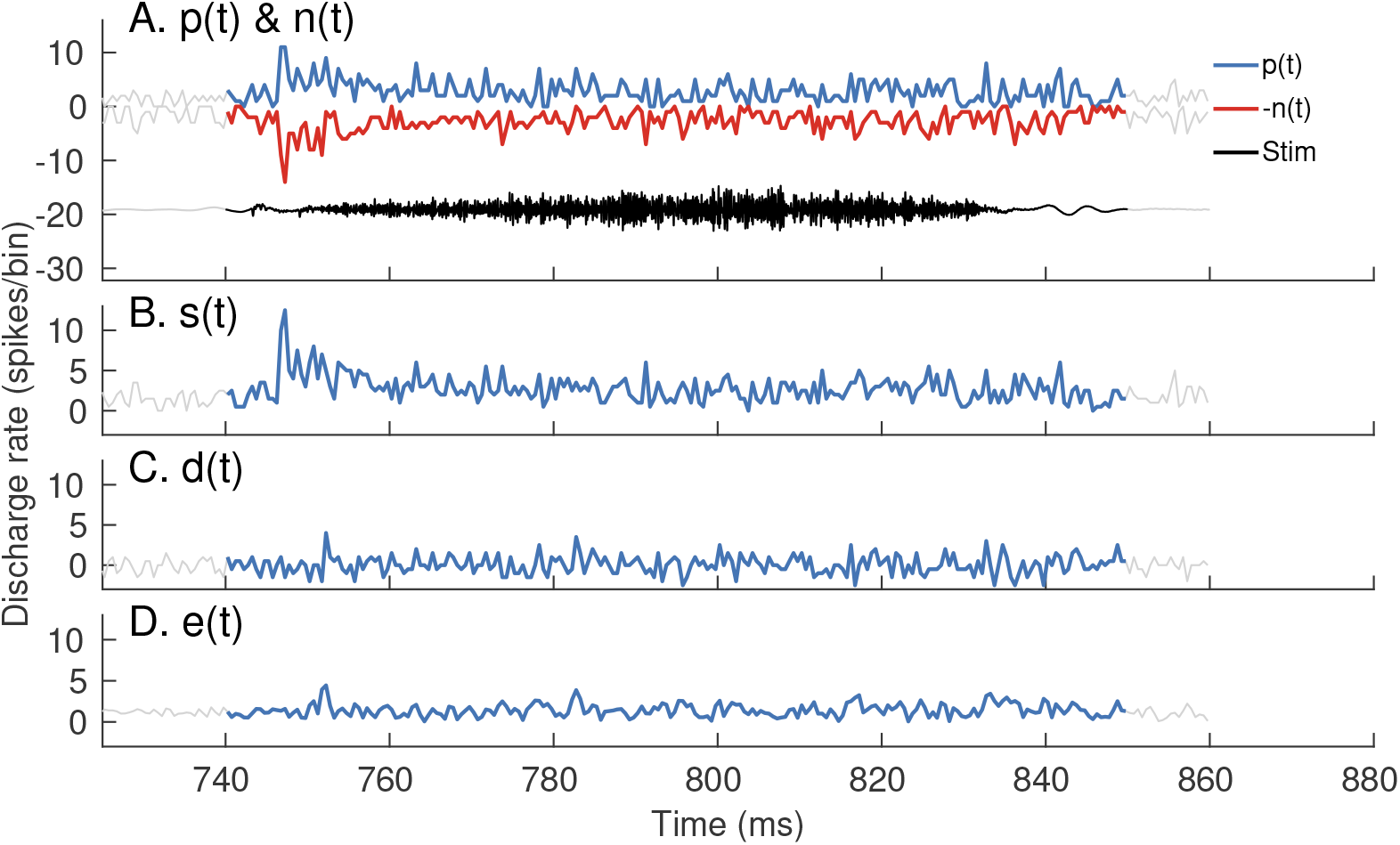
*p*(*t*), *n*(*t*), and *s*(*t*) have robust representations of the onset response, whereas *e*(*t*) and *d*(*t*) do not. Response of a high-frequency fiber (CF= 5.8 kHz, SR= 70 spikes/s) to a fricative portion (*/s/*) of the speech stimulus, *s*_3_. Stimulus intensity = 65 dB SPL. *(A)* Stimulus (black, labeled *Stim*), *p*(*t*) (blue) and *n*(*t*) (red, flipped along the y-axis). PSTH bin width = 0.5 ms. Number of stimulus repetitions per polarity = 50. (B) The sum envelope, *s*(*t*) (C) The difference PSTH, *d*(*t*), and (D) the Hilbert-envelope PSTH, *e*(*t*). Since the onset envelope is a polarity-tolerant response, all PSTHs capture the response onset except for *d*(*t*) and *e*(*t*).

Overall, these examples show that *apPSTHs* can be used to study various spectral and temporal features in neural responses for natural stimuli in the ENV/TFS dichotomy. These *apPSTHs* are summarized in Table 1 (and illustrated in S1 Fig).

### Quantifying ENV and TFS using *apPSTHs* for nonstationary signals

In the discussion so far, we have argued for using spectrally specific metrics to analyze neural responses to stationary stimuli. Another example where spectral specificity is needed is in evaluating the neural coding of nonstationary speech features (e.g., formant transitions). Speech is a nonstationary signal and conveys substantial information in its dynamic spectral trajectories (e.g., Fig 1A). A number of studies have investigated the robustness of the neural representation of dynamic spectral trajectories using frequency glides and frequency-modulated tones as the stimulus (Billings et al., 2019; Clinard and Cotter, 2015; Krishnan and Parkinson, 2000; Skoe and Kraus, 2010). These studies have usually employed a spectrogram analysis. While a spectrogram is effective for analyzing responses to nonstationary signals with unknown parameters, it does not explicitly incorporate information about the stimulus, which is often designed by the experimenter. Since the spectrogram relies on a narrow moving temporal window, it offers poor spectral resolution due to the time-frequency uncertainty principle. The same limitation applies to wavelet transforms that rely on segmenting the signal into shorter windows, even though window length varies across frequency. Instead of using these windowing-based analyses, frequency demodulation and filtering can be used together to estimate power along a spectrotemporal trajectory more accurately as described below. While this demodulation-based method has been described previously for other signals (Olhede and Walden, 2005), we apply this method to natural speech and extend this approach to construct a new spectrally compact time-frequency representation called the *harmonicgram*. These spectrally specific analyses will facilitate more sensitive metrics to investigate the coding differences between nonstationary features in natural speech and extensively studied stationary features in synthetic speech.

### Frequency-demodulation-based spectrotemporal filtering

First, we describe the spectrotemporal filtering technique using an example stimulus with dynamic spectral components (Fig 9). The 2-second long stimulus consists of three spectrotemporal trajectories: (1) a stationary tone at 1.4 kHz, (2) a stationary tone at 2 kHz, and (3) a dynamic linear chirp that moves from 400 to 800 Hz over the stimulus duration. We are interested in estimating the power of the nonstationary component, the linear chirp. In order to estimate the power of this chirp, conventional spectrograms will employ one of the following two approaches. First, one can use a long window (e.g., 2 seconds) and compute power over the 400-Hz bandwidth from 400 to 800 Hz. In the second approach, one can use moving windows that are shorter in duration (e.g., 50 ms) and compute power with a resolution of 30 Hz (20-Hz imposed by inverse of the window duration and 10-Hz imposed by change in chirp frequency over 50 ms). As an alternative to these conventional approaches, one can demodulate the spectral trajectory of the linear chirp so that the chirp is demodulated to near 0 Hz (Fig 9C and 9D, see *Materials and Methods*). Then, a low-pass filter with 0.5-Hz bandwidth (as determined by the reciprocal of the 2-s stimulus duration) can be employed to estimate the time-varying power along the chirp trajectory. This time-varying power is estimated at the stimulus sampling rate, similar to the temporal sampling of the output of a band-pass filter applied on stationary signals. While the same temporal sampling can be achieved using the spectrogram by sliding the window by one sample and estimating the chirp-related power for each window, it will be computationally much more expensive compared to the frequency-demodulation-based approach. Furthermore, the spectral resolution of 0.5 Hz is the same as that for a stationary signal, which demonstrates a 60-fold improvement compared to the 50-ms window-based spectrogram approach.

**Fig 9.**
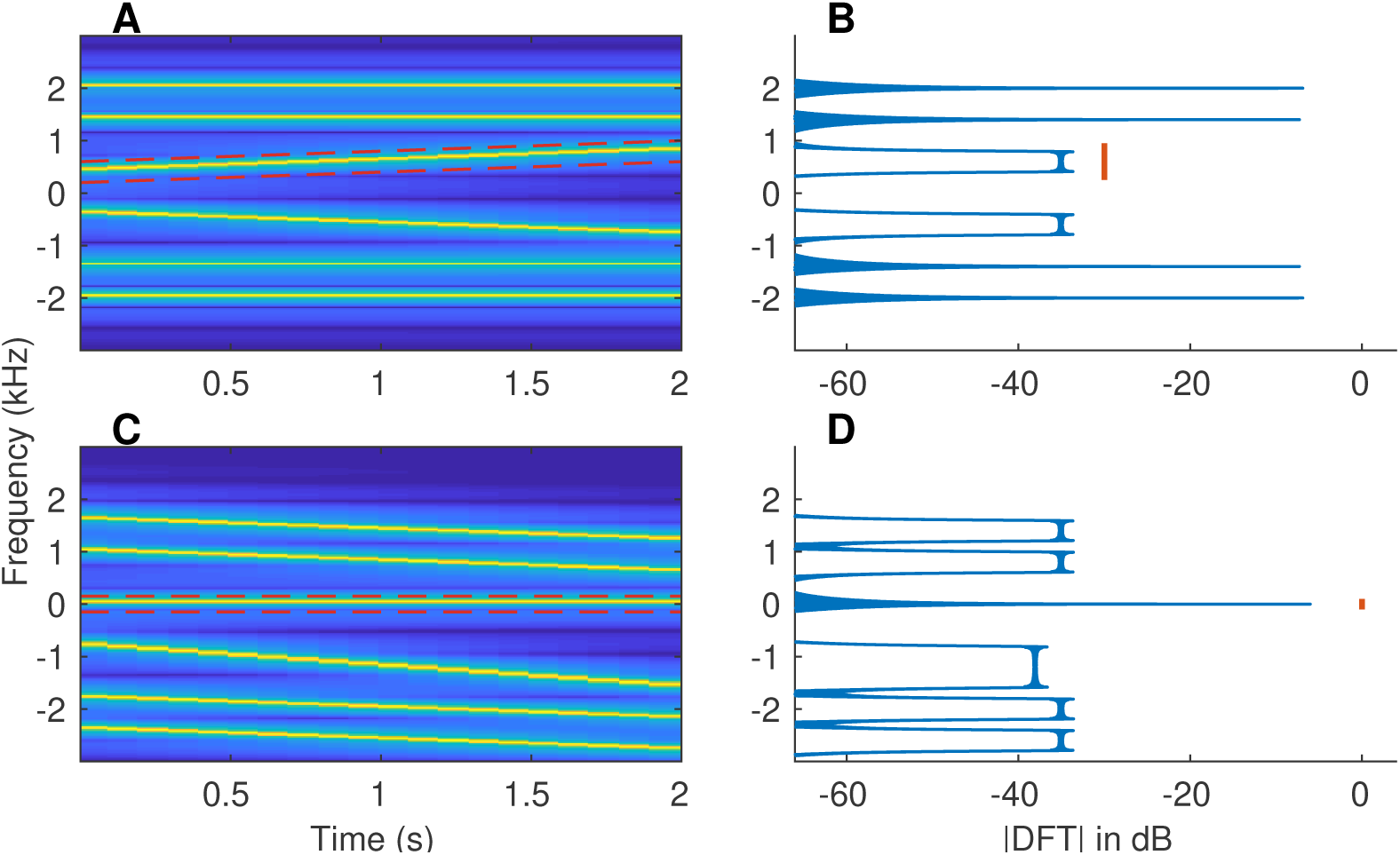
More accurate estimates of power along a spectrotemporal trajectory can be obtained using frequency demodulation. (A) Spectrogram of a synthesized example signal that mimics a single speech-formant transition. The 2-s signal consists of two stationary tones (1.4 and 2 kHz) and a linear frequency sweep (400 to 800 Hz). Red dashed lines outline the spectrotemporal trajectory along which we want to compute the power. Both positive and negative frequencies are shown for completeness. (B) Fourier-magnitude spectrum of the original signal. Energy related to the target spectrotemporal trajectory is spread over a wide frequency range (400 to 800 Hz, red line). (C) Spectrogram of the frequency-demodulated signal, where the target trajectory was used for demodulation (i.e., shifted down to 0 Hz). (D) Magnitude-DFT of the frequency-demodulated signal. The desired trajectory is now centered at 0 Hz, with its (spectral) energy spread limited only by the signal duration (i.e., equal to the inverse of signal duration), and hence, is much narrower.

### The *harmonicgram* for synthesized nonstationary speech

As shown in Fig 9, combined use of frequency demodulation and low-pass filtering can provide an alternative to the spectrogram for analyzing signals with time-varying frequency components. Such an approach can also be used to study coding of dynamic stimuli that have harmonic spectrum with time-varying *F*_0_, such as music and voiced speech. At any given time, a stimulus with a harmonic spectrum has substantial energy only at multiples of the fundamental frequency, *F*_0_, which itself can vary with time [i.e., *F*_0_(*t*)]. We take advantage of this spectral sparsity to introduce a new compact representation, the *harmonicgram*. Consider the *k* -th harmonic of *F*_0_(*t*); power along this trajectory [*kF*_0_(*t*)] can be estimated using the frequency-demodulation-based spectrotemporal filtering technique. One could estimate the time-varying power along all integer multiples (*k*) of *F*_0_(*t*). This combined representation of the time-varying power across all harmonics of *F*_0_ is the *harmonicgram* (see *Materials and Methods*). This name derives from the fact that this representation uses harmonic number instead of frequency (or spectrum) as in the conventional spectrogram.

Fig 10 shows harmonicgrams derived from *apPSTHs* in response to the nonstationary synthesized vowel, *s*_2_. The first two formants are represented by their harmonic numbers, *F*_1_(*t*)*/F*_0_(*t*) and *F*_2_(*t*)*/F*_0_(*t*), which are known a priori in this case. Two harmonicgrams were constructed using responses from two AN fiber pools: (1) AN fibers that had a low CF (CF *<* 1 kHz), and (2) AN fibers that had a medium CF (1 kHz *<* CF *<* 2.5 kHz). Previous neurophysiological studies have shown that AN fibers with CF near and slightly above a formant strongly synchronize to that formant, especially at moderate to high intensities (Delgutte and Kiang, 1984a; Young and Sachs, 1979). Therefore, the low-CF pool was expected to capture *F*_1_, which changed from 630 Hz to 570 Hz. Similarly, the medium-CF pool was expected to capture *F*_2_, which changed from 1200 Hz to 1500 Hz. The harmonicgram for each pool was constructed by using the average Hilbert-phase PSTH, *ϕ*(*t*), of all AN fibers in the pool. The harmonicgram is shown from 38 ms to 188 ms to optimize the dynamic range to visually highlight the formant transitions by ignoring the onset response. The dominant component in the neural response for *F*_1_ was expected at the harmonic number closest to *F*_1_*/F*_0_. For this stimulus, *F*_1_*/F*_0_ started at a value of 6.3 (630/100) and reached 4.75 (570/120) at 188 ms crossing 5.5 at 88.5 ms (Fig 10A). This transition of *F*_1_*/F*_0_ was faithfully represented in the harmonicgram where the dominant response switched from the 6th to the 5th harmonic near 90 ms. Similarly, *F*_2_*/F*_0_ started at 12, consistent with the dominant response at the 12th harmonic before 100 ms (Fig 10B). Towards the end of the stimulus, *F*_2_*/F*_0_ reached 12.5, which is consistent with the near-equal power in the 12th and the 13th harmonic in the harmonicgram. In contrast to findings from previous studies, the harmonicgram for the medium-CF pool indicates that these fibers respond to both the first and second formants (Delgutte and Kiang, 1984a; Miller et al., 1997). Such a complex response with components corresponding to multiple formants is likely due to the steep slope of the vowel spectrum (S4 Fig).

**Fig 10.**
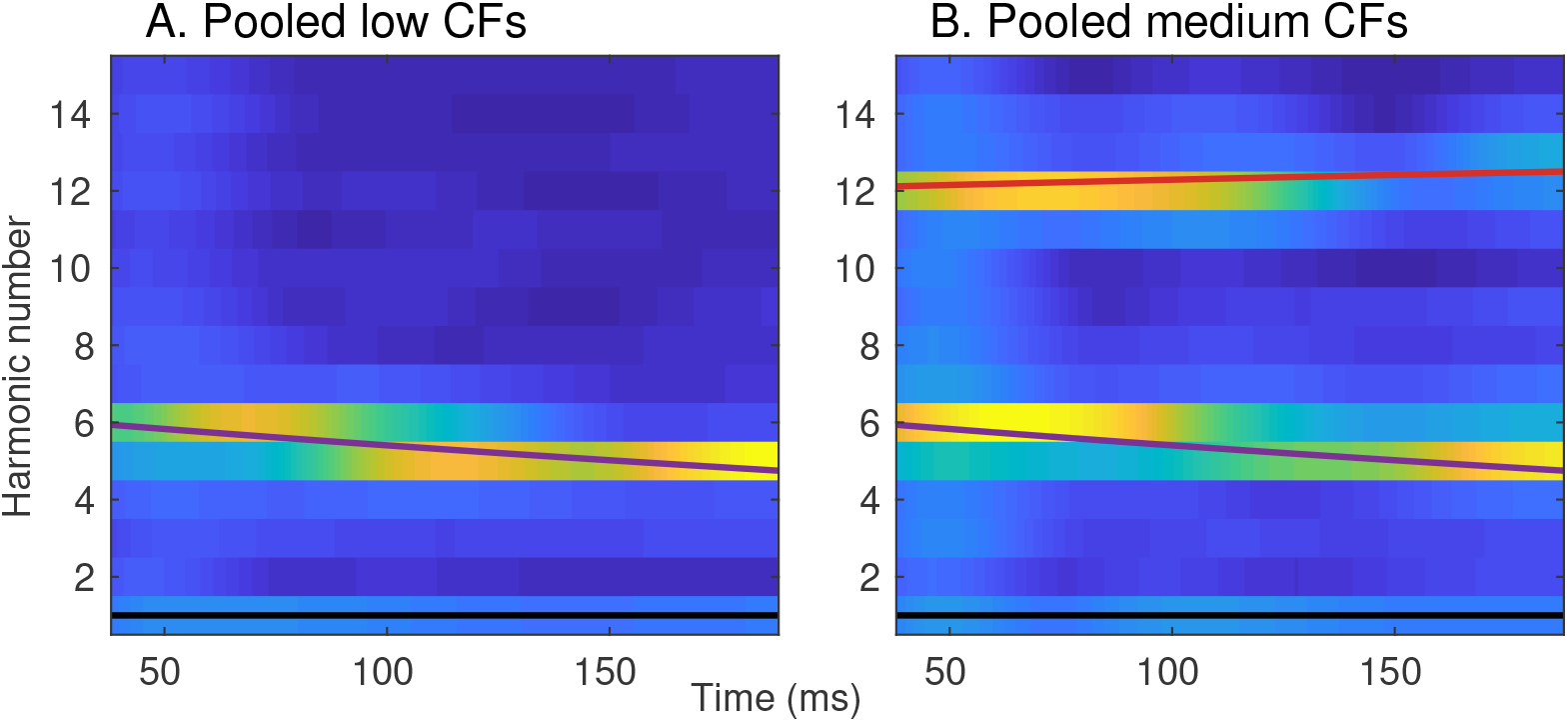
The harmonicgram can be used to visualize formant tracking in synthesized nonstationary speech. Neural harmonicgrams for fibers with a CF below 1 kHz (A, N=16) and for fibers with a CF between 1 and 2.5 kHz (B, N=29) in response to the dynamic vowel, *s*_2_. Stimulus intensity = 65 dB SPL. The formant frequencies mimic formant trajectories of a natural vowel (Hillenbrand and Nearey, 1999). A 20-Hz bandwidth was employed to low-pass filter the demodulated signal for each harmonic. The harmonicgram for each AN-fiber pool was constructed by averaging the Hilbert-phase PSTHs of all AN fibers within the pool. PSTH bin width = 50 *μ*s. Data are from one chinchilla. The black, purple and, red lines represent the fundamental frequency (*F*_0_*/F*_0_), the first formant (*F*_1_*/F*_0_) and the second formant (*F*_2_*/F*_0_) contours, respectively. The time-varying formant frequencies were normalized by the time-varying *F*_0_ to convert the spectrotemporal representation into a harmonicgram.

### The harmonicgram for natural speech

The harmonicgram analysis is not limited to synthesized vowels, but can also be applied to natural speech (Fig 11). These harmonicgrams were constructed for the natural speech stimulus, *s*_3_, using average *ϕ*(*t*) for the same low-CF and medium-CF AN fiber pools used in Fig 10. Here, we consider a 500-ms segment of the stimulus, which contains multiple phonemes. Qualitatively, similar to Fig 10, these harmonicgrams capture formant contours across phonemes. The harmonicgram for the low-CF pool emphasizes the *F*_1_ contour, whereas the harmonicgram for the medium-CF pool primarily emphasizes the *F*_2_ contour, and to a lesser extent, the *F*_1_ contour. Compared to the spectrogram, the harmonicgram representation is more compact and spectrally specific. Furthermore, from a neural-coding perspective, quantifying how individual harmonics of *F*_0_ are represented in the response is more appealing than the spectrogram since response energy is concentrated only at these *F*_0_ harmonics.

**Fig 11.**
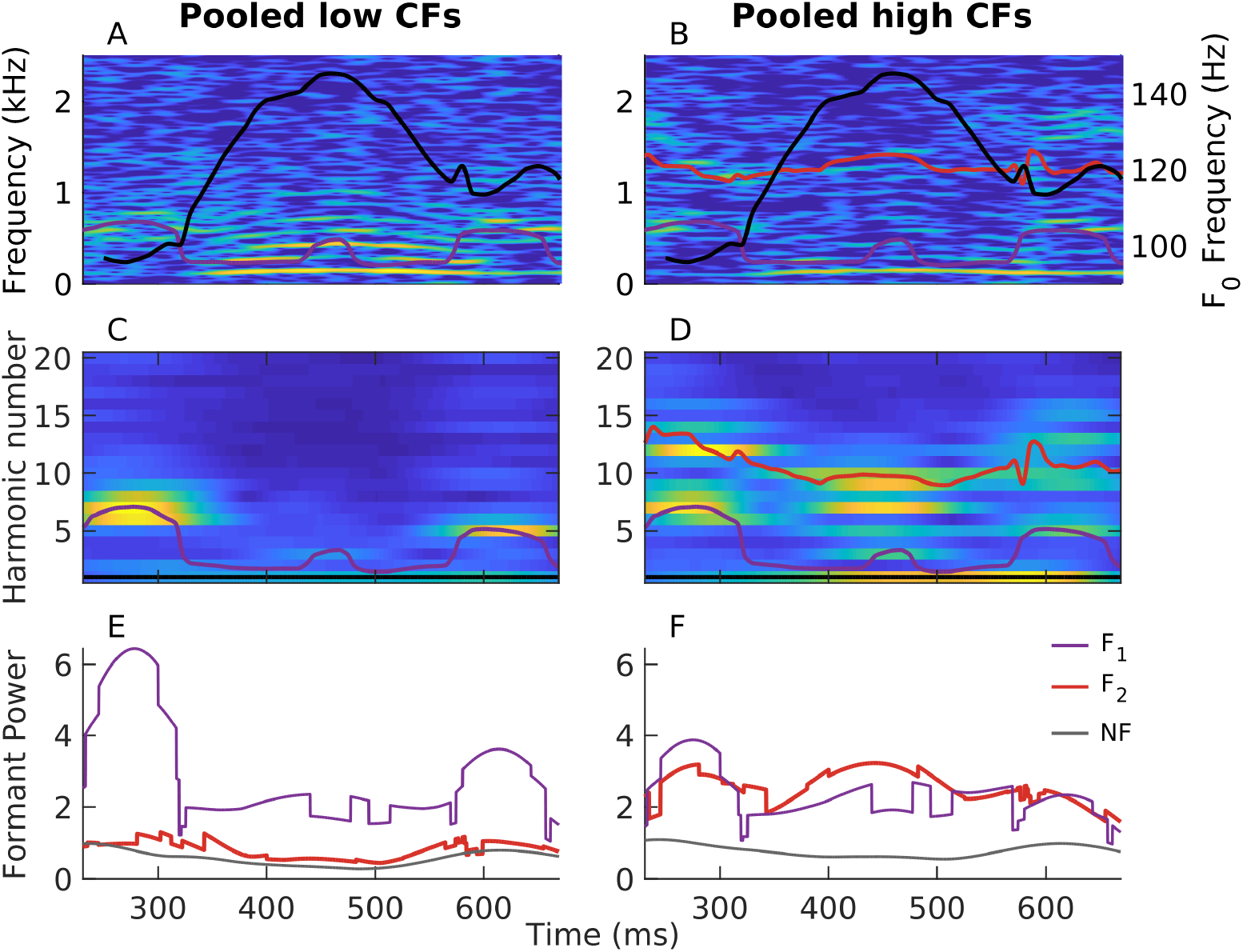
The harmonicgram can be used to quantify the coding of time-varying stimulus features at superior spectrotemporal resolution compared to the spectrogram. Harmonicgrams were constructed using *ϕ*(*t*) for the same two AN-fiber pools described in Fig 10. PSTH bin width = 50 *μ*s. A 9-Hz bandwidth was employed to low-pass filter the demodulated signal for each harmonic. The data were collected from one chinchilla in response to the speech stimulus, *s*_3_. Stimulus intensity = 65 dB SPL. A 500-ms segment corresponding to the voiced phrase “amle” was considered. (A, B) Spectrograms constructed from the average *ϕ*(*t*) for the low-CF pool (A) and from the medium-CF pool (B). (C, D) Average harmonicgrams for the same set of fibers as in A and B, respectively. Warm (cool) colors represent regions of high (low) power. The first-formant contour (*F*_1_ in A and B, *F*_1_*/F*_0_ in C and D) is highlighted in purple. The second-formant contour (*F*_2_ in A and B, *F*_2_*/F*_0_ in C and D) is highlighted in red. Trajectories of the fundamental frequency (black in A and B, right Y axis) and the formants were obtained using Praat (Boersma, 2001). (E, F) Harmonicgram power near the first formant (purple) and the second formant (red) for the low-CF pool (E) and the medium-CF pool (F). Harmonicgram power for each formant at any given time (*t*) was computed by summing the power in the three closest *F*_0_ harmonics adjacent to the normalized formant contour [e.g., *F*_1_(*t*)*/F*_0_(*t*)] at that time. The noise floor (NF) for power was estimated as the sum of power for the 29th, 30th, and 31st harmonics of *F*_0_ because the frequencies corresponding to these harmonics were well above the CFs of both fiber pools. These time-varying harmonicgram power metrics are spectrally specific to *F*_0_ harmonics and are computed with high temporal sampling rate (same as the original signal). This spectrotemporal resolution is much better than the spectrotemporal resolution that can be obtained using spectrograms.

The harmonicgram not only provides a compact representation for nonstationary signals with harmonic spectra, it can also be used to quantify coding strength of time-varying features, such as formants for speech (Figs 11E and 11F). In these examples, the strength of formant coding at each time point, *t*, was quantified as the sum of power in the three harmonics closest to the *F*_0_-normalized formant frequency at that time [e.g., *F*_1_(*t*)*/F*_0_(*t*)]. As expected, power for the harmonics near the first formant was substantially greater than for the second formant for the low-CF pool (Fig 11E). For the medium-CF pool, *F*_2_ representation was robust over the whole stimulus duration, although *F*_1_ representation was largely comparable (Fig 11F). These examples demonstrate novel analyses using the *apPSTH* -based harmonicgram to quantify time-varying stimulus features in single-unit neural responses at high spectrotemporal resolution, which is not possible with conventional windowing-based approaches.

### The harmonicgram can also be used to analyze FFRs in response to natural speech

As mentioned earlier, a major benefit of using *apPSTHs* to analyze spike trains is that the same analyses can also be applied to evoked far-field potentials. In Fig 12, the harmonicgram analysis was applied to the difference FFR recorded in response to the same speech sentence (*s*_3_) that was used in Fig 11. In fact, these FFR data and spike-train data used in Fig 11 were collected from the same chinchilla. The difference FFR was computed as the difference between FFRs to opposite polarities of the stimulus. The spectrogram and harmonicgram can also be constructed using the Hilbert-phase FFR to highlight the TFS component of the response (S5 Fig). Unlike the *apPSTHs* for AN fibers, the FFR cannot be used to construct two sets of harmonicgrams corresponding to different populations of neurons because the FFR lacks tonotopic specificity. Nevertheless, this FFR-harmonicgram is strikingly similar to the medium-CF pool harmonicgram in Fig 11D. The dynamic representations of the first two formants are robust in both the representations. In fact, the FFR representations seem more robust in formant tracking compared to PSTH-derived representations, qualitatively, especially for the harmonicgram. A more uniform sample of neurons contribute to evoked responses compared to the AN fiber sample corresponding to Fig 11, which could be a factor for the robustness of the FFR representations. Overall, these results reinforce the idea that using *apPSTHs* to analyze spike trains offers the same spectrally specific analyses that can be applied to evoked far-field potentials, e.g., the FFR, thus allowing a unifying framework to study temporal coding for both stationary and nonstationary signals in the auditory system.

**Fig 12.**
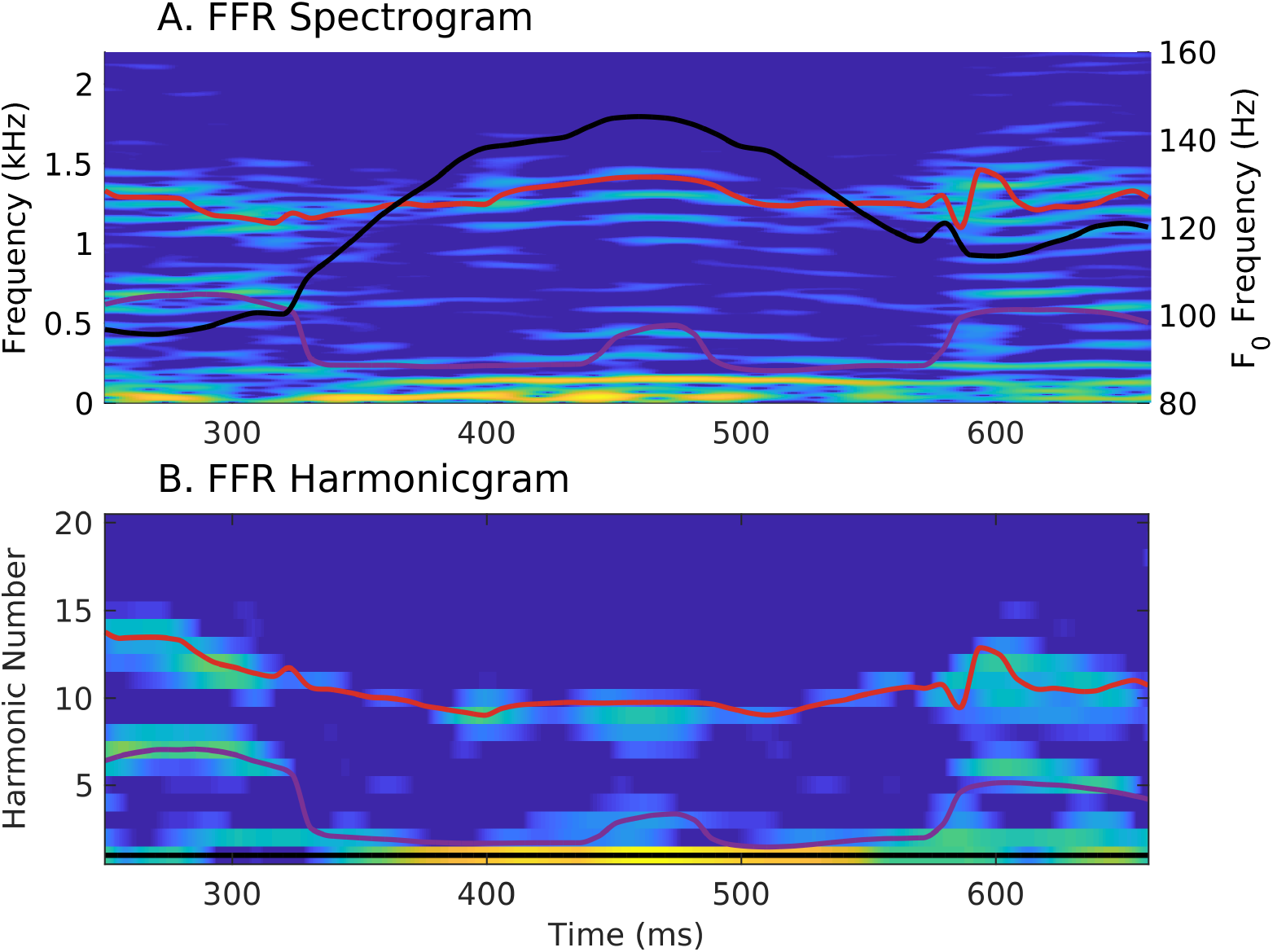
The harmonicgram of the FFR to natural speech shows robust dynamic tracking of formant trajectories, similar to the AN-fiber harmonicgram. Comparison of the spectrogram (A) and the harmonicgram (B) for the FFR recorded in response to the same stimulus, *s*_3_ that was used to analyze *apPSTHs* in Fig 11. Stimulus intensity = 65 dB SPL. Lines and colormap are the same as in Fig 11. These plots were constructed using the difference FFR, which reflects the neural coding of both stimulus TFS and ENV. To highlight the coding of stimulus TFS, Hilbert-phase [*ϕ*(*t*)] FFR can be used instead of the difference FFR (S5 Fig). The FFR harmonicgram (A) is strikingly similar to the AN-fiber harmonicgrams in Figs 11C and 11D in that the representations of the first two formants are robust. The FFR data here and spike-train data used in Fig 11 were obtained from the same animal.

## Discussion

### Use of *apPSTHs* underlies a unifying framework to study temporal coding in the auditory system

A better understanding of the neural correlates of perception requires the integration of electrophysiological, psychophysical, and neurophysiological analyses in the same framework. Although extensive literature exists in both electrophysiology and neurophysiology on the neural correlates of perception, the analyses employed in these studies have diverged. This disconnect is largely because the forms of the neural data are different (i.e., continuous-valued waveforms versus point-process spike trains). The present report provides a unifying framework for analyzing spike trains using *apPSTHs*, which offers numerous benefits over previous neurophysiological analyses. Specifically, the use of *apPSTHs* incorporates many of the previous ad-hoc approaches, such as VS and correlograms (Eqs 3 to 5). In fact, correlograms and metrics derived from them can be estimated using *apPSTHs* in a computationally efficient way. The *apPSTHs* essentially convert the naturally rectified neurophysiological point-process data into a continuous-valued signal, which allows advanced signal processing tools designed for continuous-valued signals to be applied to spike-train data. For example, *apPSTHs* can be used to derive spectrally specific TFS components [e.g., *ϕ*(*t*), Fig 6], multitaper spectra (Fig 3), modulation-domain representations (Fig 4), and harmonicgrams (Figs 10 and 11). *apPSTHs* can also be directly compared to evoked far-field responses for both stationary and nonstationary stimuli (e.g., Figs 11 and 12).

### Temporal coding metrics should be spectrally specific

The various analyses explored here advocate for spectral specificity of temporal coding metrics. The need for spectrally specific analyses arises for two reasons: (1) neural data is finite and stochastic, and (2) spike-train data are rectified. Neural stochasticity exacerbates spectral-estimate variance at all frequencies; therefore, time-domain (equivalently broadband) metrics will be noisier compared to narrowband metrics. Similarly, the rectified nature of spike-train data introduces harmonic distortions in the response spectrum, which can corrupt broadband metrics (e.g., TFS distortion at two times the carrier frequency corrupting estimates of ENV coding, Figs 5A and 5B).

These issues requiring spectral specificity are not unique to the *apPSTH* analyses but also apply to classic metrics, e.g., correlograms. For example, the broadband correlation index (CI) metric is appropriate to analyze responses of neurons with high CFs, but the CI metric is corrupted by rectifier distortions for neurons with low CFs (Heinz and Swaminathan, 2009; Joris et al., 2006). Studies have previously tried to avoid these distortions in the *sumcor* by restricting the response bandwidth to below the CF because, for a given filter, the envelope bandwidth cannot be greater than the filter bandwidth (Heinz and Swaminathan, 2009; Kale and Heinz, 2010).

Here, we have extended and generalized the analysis of these issues using narrowband stimuli. In particular, when a neuron responds to low-frequency stimulus energy that is below half the phase-locking cutoff, responses that contain any polarity-tolerant component [e.g., *p*(*t*), *n*(*t*), *s*(*t*), *SAC*, and *sumcor*] will be corrupted by rectifier distortion of the polarity-sensitive component (Fig 5E). Any broadband metric of temporal coding should exclude these distortions at twice the carrier frequency. Beyond avoiding rectifier distortion, limiting the bandwidth of a metric to only the desired bands will lead to more precise analyses by minimizing the effects of neural stochasticity (Fig 5H). For example, envelope coding metrics for SAM-tone stimuli should consider the spectrum power only at *F*_*m*_ and its harmonics (Vasilkov and Verhulst, 2019), rather than the simple approach of always low-pass filtering at CF (Heinz and Swaminathan, 2009).

Similar to envelope-based metrics, metrics that quantify TFS coding should also be spectrally specific to the carrier frequency. Previous metrics of TFS coding, such as *d*(*t*) and *difcor*, are not specific to the carrier frequency but rather include modulation sidebands as well as additional sidebands due to transduction nonlinearities (Fig 6). In contrast, *ϕ*(*t*) introduced here emphasizes the carrier and suppresses the sidebands (Fig 6). Thus, the spectrally specific *ϕ*(*t*) is a better TFS response, which relates to the zero-crossing signal used in the signal processing literature (Logan, 1977; Voelcker, 1966; Wiley, 1981).

### Spectral-estimation benefits of using *apPSTHs*

Neurophysiological studies have usually favored the DFT to estimate the response spectrum. For example, the DFT has been applied to the period histogram (Delgutte and Kiang, 1984a; Young and Sachs, 1979), the single-polarity PSTH (Carney and Geisler, 1986; Miller and Sachs, 1983), the difference PSTH (Sinex and Geisler, 1983), and correlograms (Louage et al., 2004). Since spike-train data are stochastic and usually sparse and finite, there is great scope for spectral estimates, including the DFT spectrum, to suffer from bias and variance issues. The multitaper approach optimally uses the available data to minimize the bias and variance of the spectral estimate (Babadi and Brown, 2014; Percival and Walden, 1993; Thomson, 1982). The multitaper approach can be used with both *apPSTHs* and correlograms, but using *apPSTHs* offers additional variance improvement up to a factor of 2 (Fig 3). This improvement is because twice as many tapers (both odd and even) can be used with an *apPSTH* compared to a correlogram, which is an even sequence and limits analyses to only using even tapers. Additional benefits may be achievable by combining the Lomb-Scargle approach, which is well-suited for estimating the spectrum of unevenly sampled data (e.g., spike trains), with *apPSTHs* in the multitaper framework (Springford et al., 2020).

### Benefits of spectrotemporal filtering

Analysis of neural responses to nonstationary signals has been traditionally carried out using windowing-based approaches, such as the spectrogram. Shorter windows help with tracking rapid temporal structures, but they offer poorer spectral resolution. On the other hand, larger windows allow better spectral resolution at the cost of smearing rapid dynamic features. As an alternative to windowing-based approaches, spectrotemporal filtering can improve the spectral resolution of analyses by taking advantage of stimulus parameters that are known a priori (Fig 9). This approach is particularly efficient to analyze spectrally sparse signals (i.e., signals with instantaneous line spectra, such as voiced speech). In particular, the spectral resolution is substantially improved compared to the spectrogram. In addition, while the same temporal sampling can be obtained using the spectrogram, it will be much more computationally expensive compared to the spectrotemporal filtering approach, as discussed in the following example.

The benefits of spectrotemporal filtering extend to other spectrally sparse signals, like harmonic complexes. A priori knowledge of the fundamental frequency can be used to construct the harmonicgram, which takes advantage of power concentration at harmonics of *F*_0_. This approach contrasts with the spectrogram, which computes power at all frequencies uniformly. The harmonicgram can be used to analyze both kinematic synthesized vowels (Fig 10) as well as natural speech (Fig 11). The harmonicgram is particularly useful in quantifying dominant harmonics at high temporal sampling, and is thus applicable to nonstationary signals. The harmonicgram can also be applied to evoked far-field potentials (e.g., the FFR in Fig 12). While alternatives exist to analyze spike-train data in response to time-varying stimuli (Brown et al., 2002), the present spectrotemporal technique is simpler and can be directly applied to both spike-train data and far-field responses. Overall, these results support the idea that using *apPSTHs* to analyze spike trains provides a unifying framework to study temporal coding in the auditory system across modalities. Furthermore, this framework facilitates the study of dynamic-stimulus coding by the nonlinear and time-varying auditory system.

### *apPSTHs* allow animal models of sensorineural hearing loss to be linked to psychophysical speech-intelligibility models

Speech-intelligibility models not only improve our understanding of perceptually relevant speech features, they can also be used to optimize hearing-aid and cochlear-implant strategies. However, existing SI models work well for normal-hearing listeners but have not been widely extended for hearing-impaired listeners. This gap is largely because of the fact that most SI models are based on signal-processing algorithms in the acoustic domain, where individual differences in the physiological effects of various forms of sensorineural hearing loss on speech coding are difficult to evaluate. This gap can be addressed by extending acoustic SI models to the neural spike-train domain. In particular, spike-train data obtained from preclinical animal models of sensorineural hearing loss can be used to explore the neural correlates of perceptual deficits faced by hearing-impaired listeners (Trevino et al., 2019). These insights will be crucial for developing accurate SI models for hearing-impaired listeners.

*apPSTHs* offer a straightforward means to study various speech features in the neural spike-train domain. As *apPSTHs* are in the same discrete-time continuous-valued form as acoustic signals, acoustic SI models can be directly translated to the neural domain. Many successful SI models are based on the representation of temporal envelope (Jørgensen and Dau, 2011; Relanõ-Iborra et al., 2016), although the role of TFS remains a matter of controversy (Lorenzi et al., 2006). In fact, recent studies suggest that the peripheral representation of TFS can shape central envelope representations, and thereby alter speech perception outcomes (Ding et al., 2014; Viswanathan et al., 2019). *apPSTHs* can be used to derive modulation-domain representations so that envelope based SI models can be evaluated in the neural domain (Fig 4). Similarly, the Hilbert-phase PSTH, *ϕ*(*t*), can be used to evaluate the neural representation of TFS features. These TFS results will be particularly insightful for cochlear-implant stimulation strategies that rely on the zero-crossing component of the stimulus, which closely relates to *ϕ*(*t*)(Chen and Zhang, 2011; *Grayden et al*., *2004)*.

### Translational benefits of animal models

A key motivation of this paper was to develop a framework so that insights and findings from animal models can ultimately improve our understanding of how the human auditory system processes real-life sounds, like speech. Experiments involving human subjects are typically limited to far-field responses, such as compound action potentials, frequency-following responses, and auditory brainstem responses. However, these evoked responses include contributions from multiple sources such as the cochlear microphonic, electrical interferences, and responses from several neural substrates (King et al., 2016; Verschooten and Joris, 2014); these contributions are not clearly understood. The *apPSTH* -based framework offers a straightforward way to study these contributions by comparing anatomically specific spike-train responses with clinically viable noninvasive responses.

This framework is also beneficial to develop and validate noninvasive metrics using animal models and apply these metrics to humans. For example, we demonstrated the applicability of the new spectrally compact harmonicgram approach on both spike-train data and FFR data recorded from chinchillas to evaluate speech coding. This harmonicgram analysis can also be applied to FFR data recorded from humans to study natural speech coding in both normal and impaired auditory systems. Similarly, the representation of other important response features, such as the onset and adaptation, can also be linked between invasive and noninvasive data using pre-clinical animal models of different forms of SNHL. Overall, these insights will be informative for estimating the anatomical and physiological states of humans using noninvasive measures, and how these states relate to individual differences in speech perception that currently challenge audiological rehabilitation.

### Limitations

#### Biological feasibility

The analyses proposed here aim to rigorously quantify the dichotomous ENV/TFS information in the neural response and bridge the definitions between the audio and neural spike-train domains. Methods discussed here may not all be biologically feasible. For example, the brain does not have access to both polarities of the stimulus. Thus, the PSTHs that require two polarities to be estimated, e.g., *s*(*t*), *d*(*t*), and *ϕ*(*t*), may not have an “internal representation” in the brain. This limitations also applies to correlogram metrics based on *sumcor* and *difcor*, which require two polarities of the stimulus. Thus, the use of the single-polarity PSTH [*p*(*t*)] to derive the central “internal representations” is more appropriate from a biological feasibility perspective (e.g., Fig 4). However, these various ENV/TFS components allow a thorough characterization of the processing of spectrotemporally complex signals by the nonlinear auditory system and can guide the development of more accurate speech-intelligibility models and help improve signal processing strategies for hearing-impaired listeners.

#### Alternating-polarity stimuli

Use of two polarities may not be sufficient to separate out all components underlying neural responses when more than two components contribute to neural responses at a given frequency. In particular, it may be intractable to separate out rectifier distortion when the bandwidths of ENV and TFS responses overlap. For example, consider the response of a broadly tuned AN fiber to a vowel, which has a fundamental frequency of *F*_0_. The energy at 2*F*_0_ in *S*(*f*) may reflect one or more of the following sources: (1) rectifier distortion to carrier energy at *F*_0_, (2) beating between (carrier) harmonics that are separated by 2*F*_0_, and (3) effects of transduction nonlinearities on the beating between (carrier) harmonics that are separated by *F*_0_. In these special cases, additional stimulus phase variations can be used to separate out these components (Billings and Zhang, 1994; Lucchetti et al., 2018).

#### The harmonicgram

A key drawback of applying the harmonicgram to natural speech is the requirement of knowing the *F*_0_ trajectory. *F*_0_ estimation is a difficult problem, especially in degraded speech. Thus, the harmonicgram could be inaccurate unless the *F*_0_ trajectory is known, or at least the original stimulus is known so that *F*_0_ can be estimated. A second confound is the unknown stimulus-to-response latency for different systems. Latencies for different neurons vary with their CF, stimulus frequency, and stimulus intensity. Thus, even if the acoustic spectrotemporal trajectory is known precisely, errors may accumulate if latencies are not properly accounted for. This issue will likely be minor for spectrotemporal trajectories with slow dynamics. For stimuli with faster dynamics, latency confounds can be easily minimized by estimating stimulus-to-response latency by cross-correlation and using a larger cutoff frequency for low-pass filtering.

## Materials and Methods

### Experimental procedures

Spike trains were recorded from single AN fibers of anesthetized chinchillas using standard procedures in our laboratory (Henry et al., 2019; Kale and Heinz, 2010). All procedures followed NIH-issued guidelines and were approved by Purdue Animal Care and Use Committee (Protocol No: 1111000123). Anesthesia was induced with xylazine (2 to 3 mg/kg, subcutaneous) and ketamine (30 to 40 mg/kg, intraperitoneal), and supplemented with sodium pentobarbital (∼7.5 mg/kg/hour, intraperitoneal). FFRs were recorded using subdermal electrodes in a vertical montage (mastoid to vertex with common ground near the nose) under the same ketamine/xylazine anesthesia induction protocol described above using standard procedures in our laboratory (Zhong et al., 2014). Spike times were stored with 10-*μ*s resolution. FFRs were stored with 48-kHz sampling rate. Stimulus presentation and data acquisition were controlled by custom MATLAB-based (The MathWorks, Natick, MA) software that interfaced with hardware modules from Tucker-Davis Technologies (TDT, Alachua, FL) and National Instruments (NI, Austin, TX).

### Speech stimuli

The following four stimuli were used in these experiments. (*s*_1_) Stationary vowel, ∧ (as in c<u>u</u>p): *F*_0_ was 100 Hz. The first three formants were placed at *F*_1_ = 600, *F*_2_ = 1200, and *F*_3_ = 2500 Hz. The vowel was 188 ms in duration. (*s*_2_) Nonstationary vowel, ∧: *F*_0_ increased linearly from 100 to 120 Hz over its 188-ms duration. The first two formants moved as well (*F*_1_: 630 → 570 Hz; *F*_2_: 1200 → 1500 Hz; see S4 Fig). *F*_3_ was fixed at 2500 Hz. The formant frequencies for both *s*_1_ and *s*_2_ were chosen based on natural formant contours of the vowel ∧ in American English (Hillenbrand et al., 1995; Hillenbrand and Nearey, 1999). *s*_1_ and *s*_2_ were synthesized using a MATLAB instantiation of the Klatt synthesizer (courtesy of Dr. Michael Kiefte, Dalhousie University, Canada). (*s*_3_) A naturally uttered Danish sentence [list #1, sentence #3 in the CLUE Danish speech intelligibility test, (Nielsen and Dau, 2009)]. (*s*_4_) A naturally uttered English sentence [Sentence #2, List #1 in the Harvard Corpus, (Rothauser, 1969)]. All speech and speech-like stimuli were played at an overall intensity of 60 to 65 dB SPL.

### Power along a spectro-temporal trajectory

Consider a known frequency trajectory, *f*_*traj*_(*t*), along which we need to estimate power in a signal, *x*(*t*). The phase trajectory, Φ*traj*(*t*), can be computed as

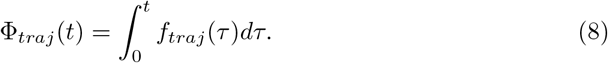

For discrete-time signals, the phase trajectory can be estimated as

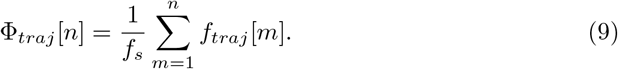

The phase trajectory can be demodulated from *x*(*t*) by multiplying a complex exponential with phase = −Φ_*traj*_(*t*) (Olhede and Walden, 2005)

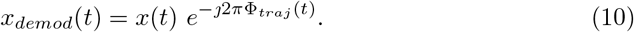

The power along *f*_*traj*_(*t*) in *x*(*t*) can be estimated as the power in *x*_*demod*_(*t*) within the spectral-resolution bandwidth (W) near 0 Hz in the spectral estimate, 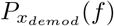.

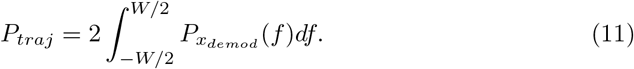

The scaling factor 2 is required because the integral in Eq 11 only represents the original positive-frequency band of the real signal, *x*(*t*); the equal amount of power within the original negative-frequency band, which is shifted further away from 0 Hz by Φ_*traj*_(*t*), should also be included (see Fig 9).

### The harmonicgram

Consider a harmonic complex, *x*(*t*), with a time-varying (instantaneous) fundamental frequency, *F*_0_(*t*). For a well-behaved and smooth *F*_0_(*t*), energy in *x*(*t*) will be concentrated at multiples of the instantaneous fundamental frequency, i.e., *kF*_0_(*t*). Thus, *x*(*t*) can be represented by the energy distributed across the harmonics of the fundamental. The time-varying power along the *k*-th harmonic of *F*_0_(*t*) can be estimated by first demodulating *x*(*t*) with the *kF*_0_(*t*) trajectory using Eq 10, and then using an appropriate low-pass filter to limit energy near 0 Hz (say within ±*W/*2). We define the *harmonicgram* as the matrix of time-varying power along all harmonics of the fundamental frequency. Thus, the harmonicgram is

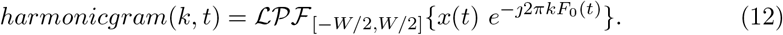

## Acknowledgments

This work was supported by an International Project Grant from Action on Hearing Loss (UK) and by NIH/NIDCD (R01-DC009838), both awarded to MGH. We thank Keith Kluender for his help with stationary and kinematic vowel synthesis. We also thank François Deloche, Hannah Ginsberg, Caitlin Heffner, and Vibha Viswanathan for their valuable feedback on an earlier version of this manuscript.

## Supporting information

**S1 Text**. Classic metrics for quantifying temporal coding in the auditory system.

**S1 Appendix**. Vector strength metric definitions.

**S2 Appendix**. Relation between the *vector strength* metric and the *difference* PSTH.

**S3 Appendix**. Relation between *shuffled correlograms* and *apPSTHs*.

**S4 Appendix**. Relation between *difcor/sumcor* and *difference/sum PSTHs*.

**S5 Appendix**. Relation between *shuffled-correlogram* peak-height and *apPSTHs*.

****S1** Table**. Glossary of terms and definitions.

****S2** Table**. Parameters for the AN model.

**S1 Fig**. Graphical illustration of *apPSTHs* in Table 1

**S2 Fig**. Nonlinear inner-hair-cell transduction function introduces additional sidebands in the spectrum for a SAM tone.

**S3 Fig**. Neural characterization of ENV and TFS using *apPSTHs* for a synthesized stationary vowel.

**S4 Fig**. DFT-magnitude for the nonstationary vowel, *s*_2_.

**S5 Fig**. FFR harmonicgram can be constructed using the Hilbert-phase response.

## Supporting information

### S1 Text. Classic metrics for quantifying temporal coding in the auditory system

Various approaches and metrics have been developed to quantify auditory temporal coding in neurophysiological responses. In this section, we motivate the need for a unified framework for auditory temporal coding by briefly reviewing these classic metrics and discussing their benefits and limitations.

### Period-histogram based metrics

The ability of AN fibers to follow the temporal structure of an acoustic stimulus has been known for a long time (Galambos and Davis, 1943). Using tones as stimuli, Kiang and colleagues showed that AN fibers prefer to discharge spikes around a particular phase of the stimulus cycle (Kiang et al., 1965). Their analysis was qualitative and involved the period histogram, which is constructed as the histogram of spike times modulo the period of one stimulus cycle (e.g., Fig 1D). Rose and colleagues used the period histogram to quantify the preference of neurons to fire during one half-cycle of a periodic stimulus (Rose et al., 1967). They introduced a metric, called the coefficient of synchronization, which is defined as the ratio of the spike count in the most effective half-cycle to the spike count during the whole stimulus cycle. The coefficient of synchronization ranges from 0.5 (for a flat period histogram) to 1.0 (for all spikes within one half-cycle). The coefficient of synchronization does not truly quantify the strength of phase locking to the stimulus cycle as it does not consider the spread of the period histogram. For example, two period histograms, one where all spikes occur at the peak of the stimulus cycle (strong phase locking), and the other where all spikes are uniformly distributed across one stimulus half-cycle (weak phase locking), will yield the same coefficient of synchronization of 1.0.

A more sensitive measure of phase locking derived from the period histogram is the vector strength [VS (Goldberg and Brown, 1969; Greenwood and Durand, 1955)], which is identical to the synchronization index metric described by Johnson (Johnson, 1980). VS has been used extensively to quantify phase-locking strength in spike-train recordings in response to periodic stimuli (Joris et al., 2004; Palmer and Russell, 1986), including stationary speech (Young and Sachs, 1979). In this framework, each spike is treated as a complex vector that has a magnitude of 1 and an angle that is defined by the spike phase relative to the stimulus phase; VS is defined as the magnitude of the average of all such vectors for spikes pooled across all stimulus repetitions (S1 Appendix). VS is a biased estimator of the “true” vector strength (Mardia, 1972) and can reach spuriously high values at low spike counts (Yin et al., 2010). This problem is avoided by using a modification of the vector strength, called the phase-projected vector strength (*V S*_*pp*_) (Yin et al., 2010). Similar approaches have been used in electrophysiological studies (Vinck et al., 2011). *V S*_*pp*_ differs from *V S* in that trial-to-trial phase consistency is also considered in computing *V S*_*pp*_ (S1 Appendix).

Overall, the period histogram and metrics derived from it (*V S* and *V S*_*pp*_) work well for applications involving stationary signals with periodic TFS (e.g., tones, Fig. 1D), ENV (e.g., sinusoidally amplitude-modulated noise), or both (e.g., sinusoidally amplitude-modulated tones, Fig. 1F). However, the period histogram ignores nonstationary features in the response that arise from the auditory system. For example, spikes in the first few stimulus cycles are often ignored while constructing the period histogram to avoid the nonstationary onset response. Similarly, since spikes corresponding to different stimulus cycles are wrapped onto a single cycle, effects of adaptation are not captured in the period histogram. Moreover, its application to nonstationary or aperiodic stimuli (e.g., natural speech) is not straightforward.

### Peristimulus-time-histogram (PSTH) based metrics

The single-polarity PSTH, *p*(*t*), is constructed as the histogram of spike times pooled across all stimulus repetitions at a certain bin width (e.g., Fig 1C). As the PSTH shows the rate variation along the course of the stimulus, it captures the onset as well as adaption effects in the response (Kiang et al., 1965; Westerman and Smith, 1988). *p*(*t*) has been applied to analyze spike trains recorded in response to periodic signals, both in the temporal and spectral domains (Delgutte, 1980; Palmer et al., 1986; Young and Sachs, 1979). A limitation of the *p(t)*-spectrum is that it is corrupted by harmonic distortions due to the rectified nature of the PSTH response (Young and Sachs, 1979). For example, the spectrum of a PSTH constructed using spike trains recorded from an AN fiber in response to a tone (*F*_*c*_) can show energy at *F*_*c*_ as well as 2*F*_*c*_ even though the stimulus itself does not have energy at 2*F*_*c*_. These issues related to rectifier distortion can be minimized by using both polarities of the stimulus (S1 Fig). Similar to the period histogram, the PSTH can also be used to derive phase-locking metrics, such as *V S* and *V S*_*pp*_. These synchrony-based metrics have been recently overshadowed by correlogram-based metrics, which are described next, since the synchrony-based metrics are limited to periodic signals. In contrast, correlogram-based approaches offer more general metrics to evaluate temporal coding of both periodic and aperiodic stimuli in the ENV/TFS dichotomy.

### Interspike-interval (ISI) based approaches (e.**g**., **correlograms)**

Interspike interval histogram analyses were developed to quantify the correlation between two spike trains, either from the same neuron or from different neurons (Hagiwara, 1954; Perkel et al., 1967a,b; Rodieck et al., 1962). Interspike intervals between adjacent spikes (also called first-order intervals) within a stimulus trial are used to construct per-trial estimates of the ISI histogram, which are then averaged across trials to form the final first-order ISI histogram (Fig ST1-C). An alternative to the first-order ISI histogram, called the all-order ISI histogram (or the autocorrelogram), can be estimated in a similar way with the only difference being the inclusion of intervals between all spikes within a trial (not only adjacent spikes) to construct the histogram (Fig ST1-E) (Møller, 1970; Rodieck, 1967). The autocorrelogram has been used to study the temporal representation of stationary as well as nonstationary stimuli (Bourk, 1976; Cariani and Delgutte, 1996a,b; Sinex and Geisler, 1981). While the autocorrelogram is attractive for its simplicity, it is confounded by refractory effects (Figs ST1-E and ST1-F). In particular, since successive spikes within a single trial cannot occur within the refractory period, the autocorrelogram shows an artifactual absence of intervals for delays less than the 0.6-ms refractory period (Fig ST1-E). As a result, the autocorrelogram spectrum is partly corrupted.

Joris and colleagues extended these ISI-based analyses to remove the confounds of the refractory effects by including all-order interspike intervals *across* stimulus trials to compute a shuffled correlogram (Louage et al., 2004). A shuffled correlogram computed using spike trains in response to multiple repetitions of a single stimulus from a single neuron is called the shuffled autocorrelogram (or the *SAC*, Fig ST1-G). Similarly, a shuffled correlogram computed using spike trains from different neurons, or for different stimuli, is called the shuffled cross-correlogram (or the *SCC*). The use of across-trial all-order ISIs provides substantially more smoothing than simple all-order ISIs because many more intervals are included in the histogram (compare Fig ST1-E with Fig ST1-G, and Fig ST1-F with Fig ST1-H).

In addition, both polarities of the stimulus can be used to separate out ENV and TFS components from the response. Stimuli with alternating polarities share the same envelope, but their phases (TFS) differ by a half-cycle at all frequencies. By averaging shuffled autocorrelograms for both stimulus polarities and shuffled cross-correlograms for opposite stimulus polarities, the *polarity-tolerant* (ENV) correlogram (called the *sumcor*) is obtained (Louage et al., 2004) (S4 Appendix). Similarly, the *polarity-sensitive* (TFS) correlogram, the *difcor*, is estimated as the difference between the average autocorrelogram for both stimulus polarities and the cross-correlogram for opposite stimulus polarities (S4 Appendix). These functions have been preferred over PSTH-based analyses for estimating correlation sequences and response spectra (Cedolin and Delgutte, 2005; Joris et al., 2006; Rallapalli and Heinz, 2016). Shuffled autocorrelograms have also been used to derive temporal metrics, such as the *correlogram peak-height* and *half-width*, to quantify the strength and precision of temporal coding in the response, respectively (Louage et al., 2004), including for nonstationary stimuli (Paraouty et al., 2018; Sayles et al., 2015; Sayles and Winter, 2008). In addition, cross-correlograms have been used to develop metrics to quantify ENV/TFS similarity between responses to different stimuli recorded from the same neuron [e.g., speech stimuli (Heinz and Swaminathan, 2009; Rallapalli and Heinz, 2016; Swaminathan and Heinz, 2012)], or between responses from different neurons (Heinz et al., 2010; Joris et al., 2006; Swaminathan and Heinz, 2011).

Although correlogram-based analyses provide a rich set of temporal metrics, they suffer from three major limitations. First, correlograms discard phase information in the response. Response phase can convey important information, especially for complex stimuli, like speech (Delgutte et al., 1998; Greenberg and Arai, 2001; Paliwal and Alsteris, 2003). Second, metrics derived from the shuffled autocorrelogram and the *sumcor* are corrupted by rectifier distortions (e.g., Fig ST1-H). Third, spectral estimates based on correlograms are appropriate for second-order stationary signals. To accommodate for nonstationary signals, usually a sliding-window-based approach is employed where in each temporal window the spectrum and/or correlogram is computed (Sayles and Winter, 2008). This windowing-based approach faces the classic problem of a time-frequency resolution trade-off. In addition, the smoothing benefit provided by the correlogram comes at large computational cost as its computation requires all-order spike-time differences across all trials. This computation cost scales quadratically (*N* ^2^) with the number of spikes (*N*) and can be cumbersome for large *N*.

## S1 Appendix. Vector strength metric definitions

### Vector Strength

The *vector strength* (*VS*) metric is used to quantify how well spikes in a spike train are synchronized to a frequency, *f* (Goldberg and Brown, 1969; Johnson, 1980). Let us denote a spike train with N spikes as <u>ζ</u> such that *<u>ζ</u>* = {*t*_1_, *t*_2_, …, *t*_*N*_} and the {*t*_*i*_}s are individual spike times. To compute the vector strength, these spike times are first transformed onto the unit circle such that *t*_*i*_ maps to *z*_*i*_ as

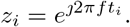

The mean of the set of complex vectors corresponding to all N spikes is

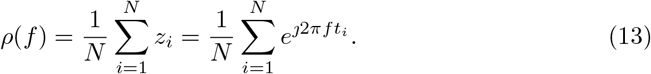

Then, *VS* at frequency *f* is defined as the magnitude of *ρ*(*f*).

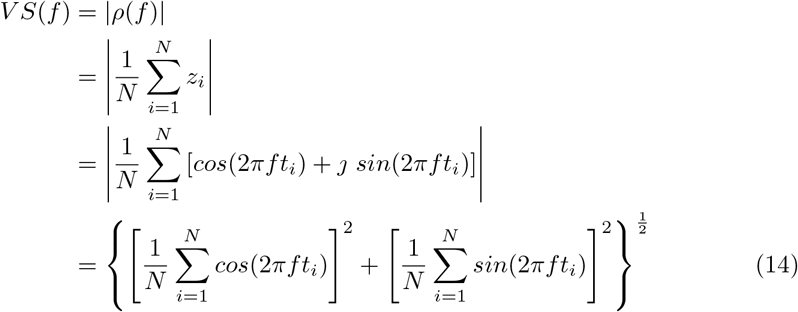

### Phase-projected Vector Strength

The *phase-projected vector strength* (*V S*_*pp*_) is identical to the *VS* for a single spike train (i.e., for a single stimulus repetition), but these metrics differ when multiple (*R*) stimulus repetitions are used. *V S*_*pp*_ is advantageous relative to *V S* when there are relatively fewer spikes per repetition (Yin et al., 2010). To estimate *V S*_*pp*_ at frequency *f*, the magnitude (i.e., *V S*) and phase [*ϕ*_*r*_(*f*)] of the mean complex vector are first calculated for individual repetitions using Eqs 13 and 14 (instead of pooling spike times across all *R* repetitions). The per-repetition VS estimates, called *V S*^*r*^(*f*), are weighted by the cosine of the phase difference between *ϕ*^*r*^(*f*) of the repetition and the mean phase based on all spikes from all repetitions, *ϕ*^*ref*^ (*f*), to estimate the *phase-projected vector strength*, 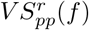, for the repetition.

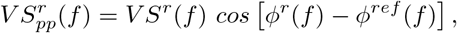

where *ϕ*^*r*^(*f*) for repetition r with *N*_*r*_ spikes 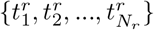 is computed as

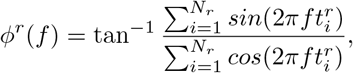

and *ϕ*^*ref*^ (*f*) is computed using all spikes across all R repetitions as

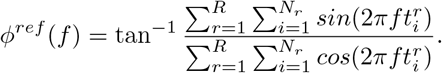

*V S*_*pp*_(*f*) for R repetitions is computed as the mean 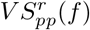 across all repetitions,

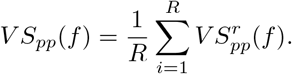

## S2 Appendix. Relation between the *vector strength* metric and the *difference PSTH*

Let us assume that we have *R* sets of spike trains {*<u>ζ</u>i*} : *i* ∈ [1, *…, R*] for a tone stimulus with duration *D* and frequency *f*_0_. Let the corresponding PSTH be *p*(*t*), and the total number of spikes be *N*.

In Eq 13, 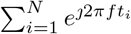 can be written as (van Hemmen, 2013)

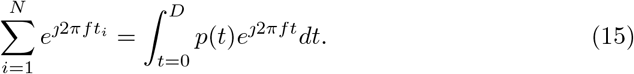

Using Eq 15 in Eq 13, we get

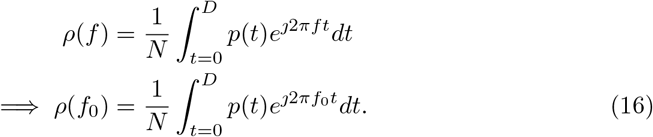

If we assume response phase locking to positive and negative polarity of a sinusoid (*f*_0_) differ by a phase of *π* [i.e., a time difference of *T*_0_*/*2(= 1*/*2*f*_0_) such that 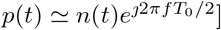, we can write

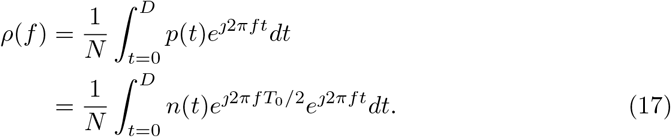

For *f* ≠ *f*_0_, the integral in Eq 17 will be zero. For *f* = *f*_0_,

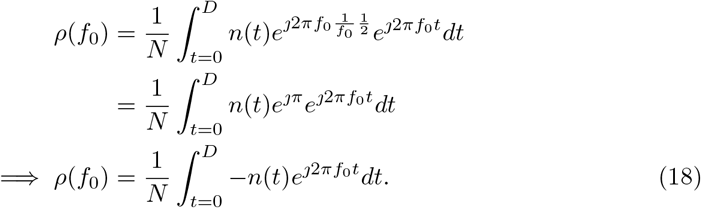

Adding Eqs. 16 and 18, we get

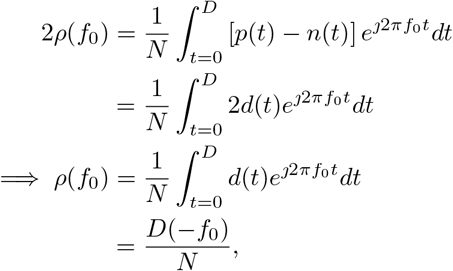

where 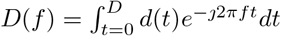 is the Fourier transform of *d*(*t*). Since *d*(*t*) is a real signal, |*D*(*f*) | = | *D*(*f*) |. Thus, the relation between VS and the difference PSTH becomes,

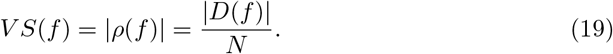

### S3 Appendix. Relation between *shuffled correlograms* and *apPSTHs*

Consider 𝕏: a set of *T*_*X*_ spike trains 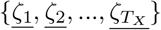 in response to a stimulus of duration *D*. For each spike train *<u>ζ</u>*_*<u>i</u>*_, we can construct a PSTH, *<u>x</u>*_*<u>i</u>*_, with PSTH bin width Δ so that the length of the single-trial PSTH *<u>x</u>*_*<u>i</u>*_ is *M* = *D/*Δ. The single-trial PSTH is a binary-valued vector because each element in the vector is either 0 or 1. Let us denote the PSTH for 𝕏 by *PSTH*_*X*_ such that 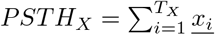. Consider 𝕐: another set of *T*_*Y*_ spike trains, with *<u>y</u>*_*<u>i</u>*_ and *PSTH*_*Y*_ defined similarly to *x*_*i*_ and *PSTH*_*X*_, respectively. Let us assume that the stimulus duration and bin width for *y*_*i*_ are the same as that for *<u>x</u>*_*<u>i</u>*_. Let the average discharge rates (in spikes/s) for 𝕏 and 𝕐 be *r*_*X*_ and *r*_*Y*_, respectively. The shuffled cross-correlogram (*SCC*) for two spike trains *<u>ζ</u>*_*<u>i</u>*_ and *<u>ζ</u>*_*<u>j</u>*_ computed using tallying (Louage et al., 2004) is identical to the cross-correlation function (denoted by *ℛ* _*𝒳𝒴*_) between their respective PSTHs, (*<u>x</u>*_*<u>i</u>*_ and *<u>x</u>*_*<u>j</u>*_). Thus, the raw (not normalized) shuffled cross-correlogram (*SCC*^*raw*^) at *τ* delay can be computed as

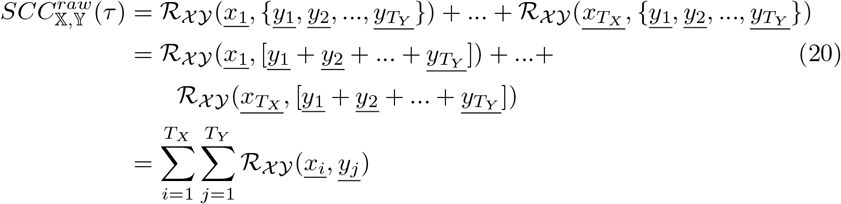

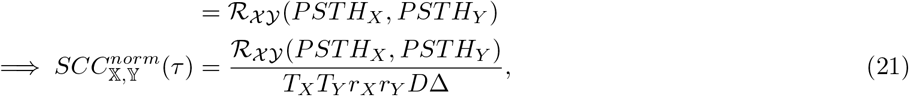

where *SCC*^*norm*^ is the normalized SCC (Heinz and Swaminathan, 2009; Louage et al., 2004).

Similarly, the raw shuffled autocorrelogram (*SAC*^*raw*^) at *τ* delay can be computed as,

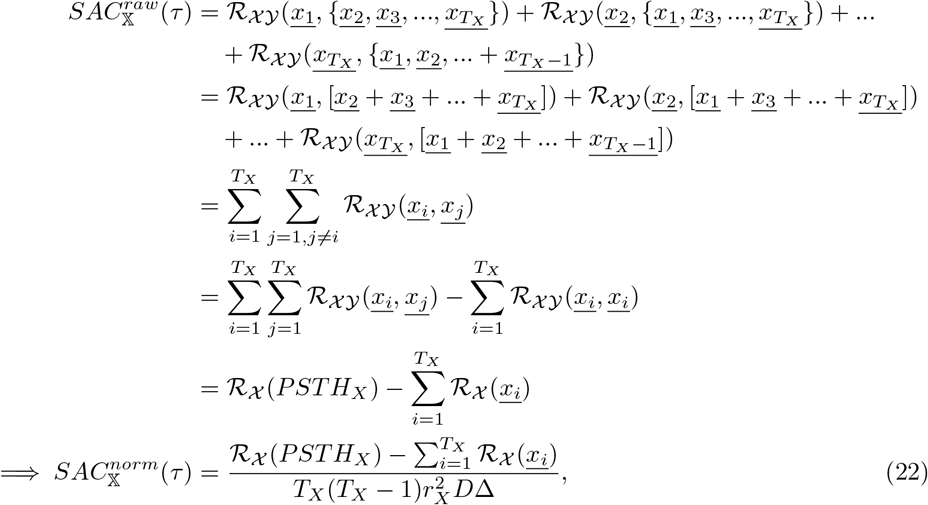

where *ℛ* _*𝒳*_ denotes the autocorrelation function. Similar to autocorrelation functions, the *SAC*^*norm*^ has its maximum at zero delay.

In the numerator of Eq 22, the term 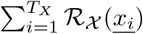 is negligible compared to *ℛ* _*𝒳*_ (*PSTH*_*X*_) for *τ* ≠0. For 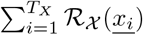 is equal to the total number of spikes (*N*) in 𝕏. Thus, Eq 22 can be further approximated by,

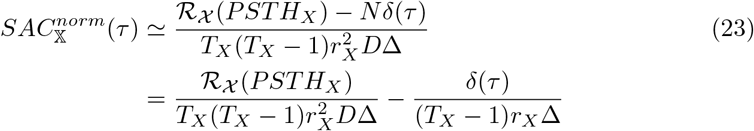

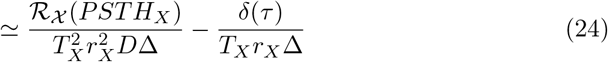

where *N* = *r*_*X*_*DT*_*X*_, and *δ* is the Dirac delta function. The simplifying approximation in Eq 24 is valid for typically used *T*_*X*_ values in neurophysiological experiments, and equates the normalization factors between *SACs* and *SCCs* when working with *difcor* and *sumcor* (e.g., S4 Appendix). Eqs 21 to 24 indicate that correlograms can be computed much more efficiently using *apPSTHs* instead of by tallying spike times [*𝒪* (*N*) instead of *𝒪* (*N* ^2^), see main text].

## S4 Appendix. Relation between *difcor/sumcor* and *difference/sum PSTHs*

Consider 𝕏_+_: spike trains in response to the positive polarity of a stimulus, and 𝕏_−_: spike trains in response to the negative polarity of the stimulus. Then, the *difcor* at *τ* delay can be computed as

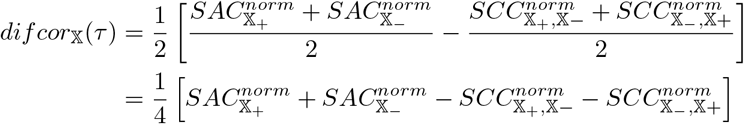

For analytic simplicity, we use Eq 24 for *SAC*^*norm*^ instead of Eq 22. Let us assume that the number of repetitions and average rates for both polarities are the same. Thus,

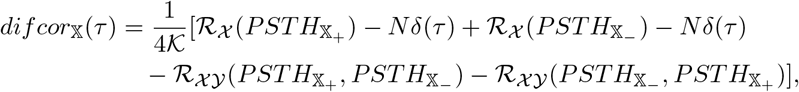

where 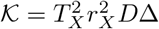 is a constant. Now, 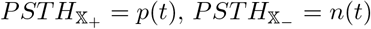, and the difference PSTH *d*(*t*) = [*p*(*t*) − *n*(*t*)] */*2. Then, the *difcor* for X at delay *τ* is

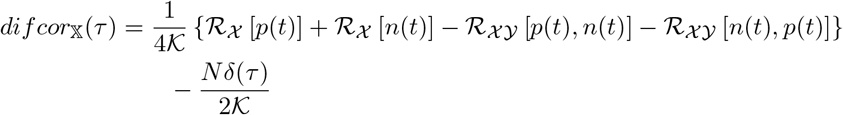

Now,

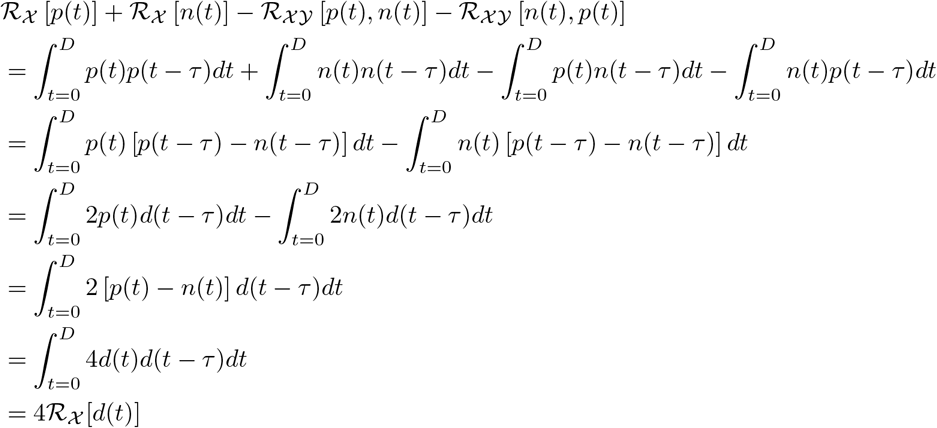

Thus,

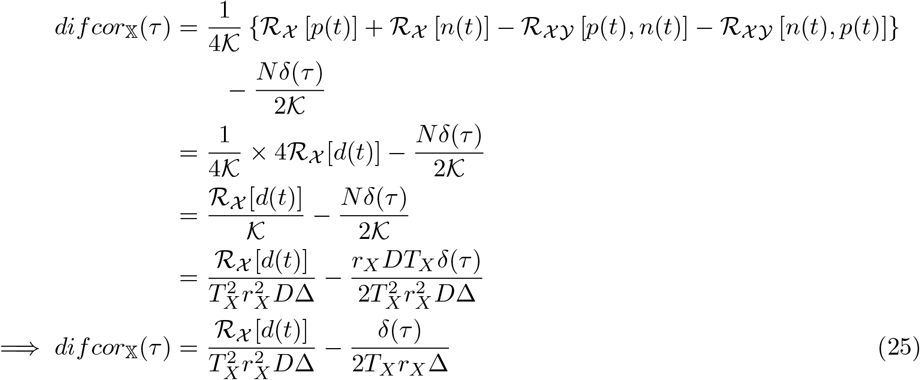

Similarly, it can be shown that

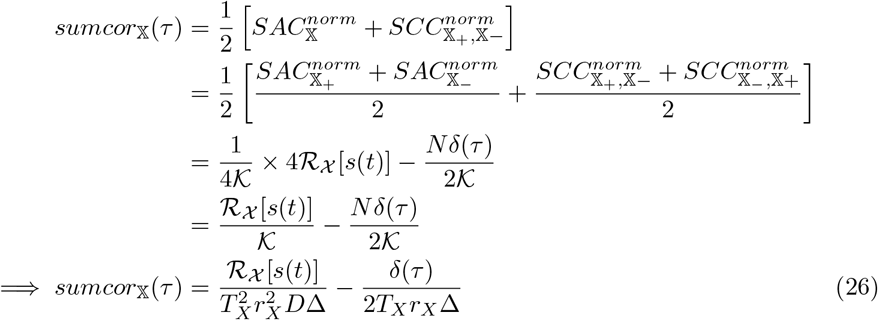

where *s*(*t*) is the sum PSTH, i.e., *s*(*t*) = [*p*(*t*) + *n*(*t*)]*/*2.

Eqs 25 and 26 indicate that *sumcor* and *difcor* are related to the autocorrelation function of the *sum* and *difference* PSTHs, respectively, and thus can be computed much more efficiently [*𝒪* (*N*) rather than *𝒪* (*N* ^2^)].

## S5 Appendix. Relation between *shuffled-correlogram* peak-height and *apPSTHs*

Consider a difference PSTH, *d*(*t*), based on a set of spike trains 𝕏 in response to a stimulus of duration *D*. Let us denote the Fourier transform of *d*(*t*) by *D*(*f*). Then, from Eq 25, the *difcor* peak-height, i.e., *difcor* value at zero delay (*τ*), can be computed as

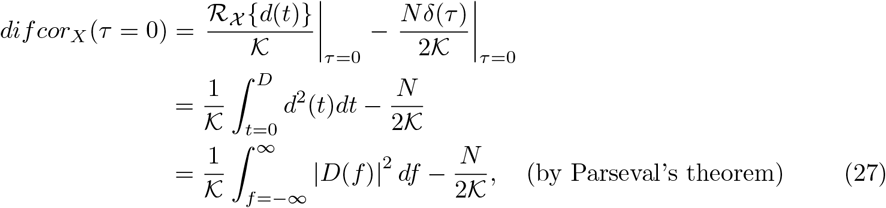

Following similar steps from Eq 26, it can also be shown that the *sumcor* peak-height can be computed as

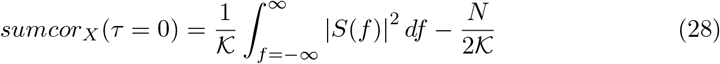

where *S*(*f*) is the Fourier transform of the sum PSTH, *s*(*t*).

Comparing Eq 19 with Eqs. 27 and 28, we see that vector strength is a frequency-specific metric, whereas correlogram peak-heights are broadband measures, which are thus susceptible to rectifier distortion (see Fig 5).

**S1 Table.**
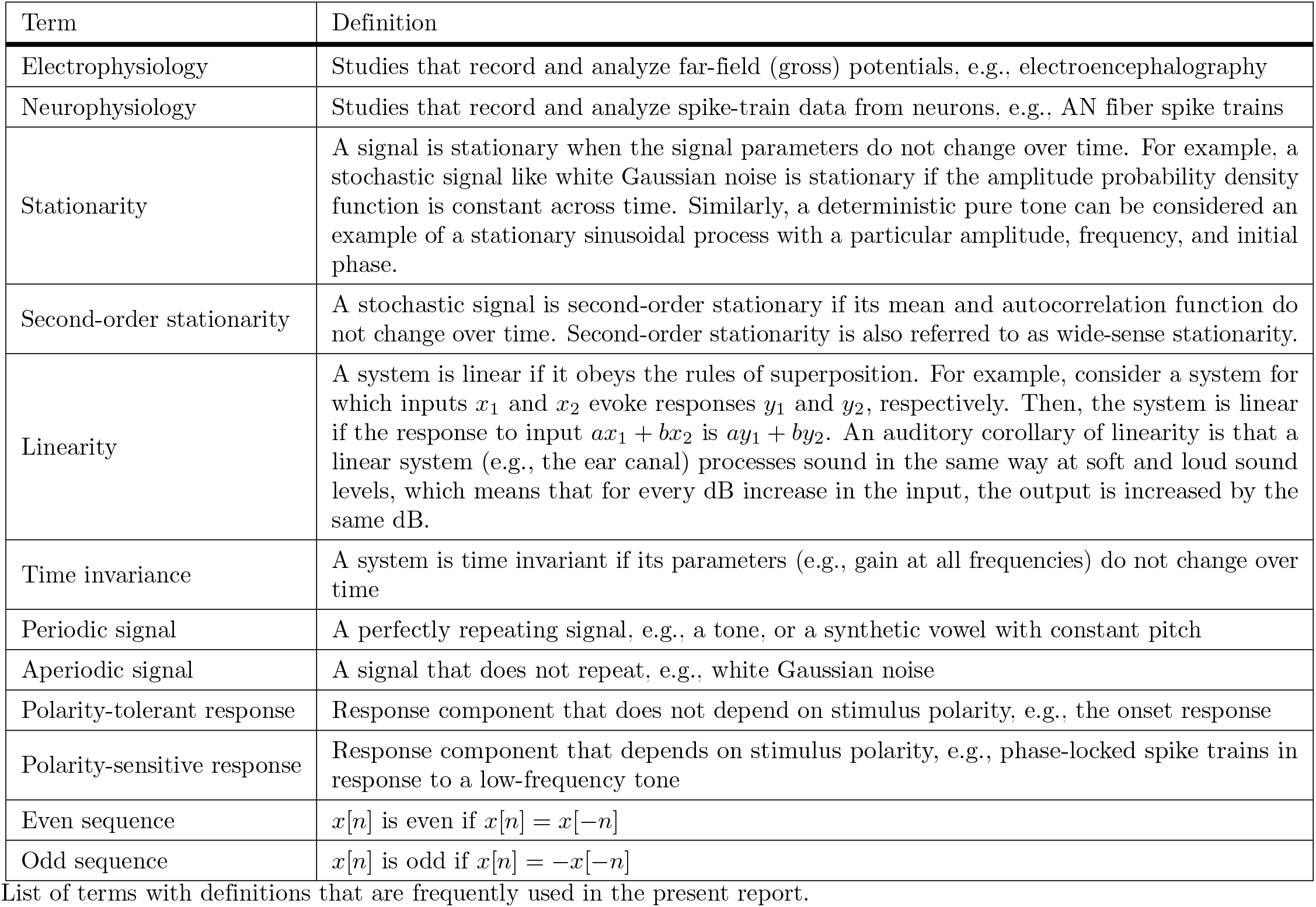
Glossary of terms and definitions.

**S2 Table.** AN model parameters.

| Parameter | Value |
| --- | --- |
| Sampling Frequency | 100 kHz |
| Number of Repetitions (per polarity) | 25 |
| Spontaneous firing rate (SR) | 70 spikes/s |
| Absolute refractory period | 0.6 ms |
| Baseline mean relative refractory period | 0.6 ms |
| OHC health value | 1.0 (normal) |
| IHC health value | 1.0 (normal) |
| Species | 1 (cat) |
| Fractional Gaussian noise type | 0 (fixed) |
| Implementation type of the power-law functions in the Synapse | 0 (approximate) |
| Spike time resolution | 10 μs |

List of parameters used in the AN model to generate simulated spike-train data.

**Fig ST1.**
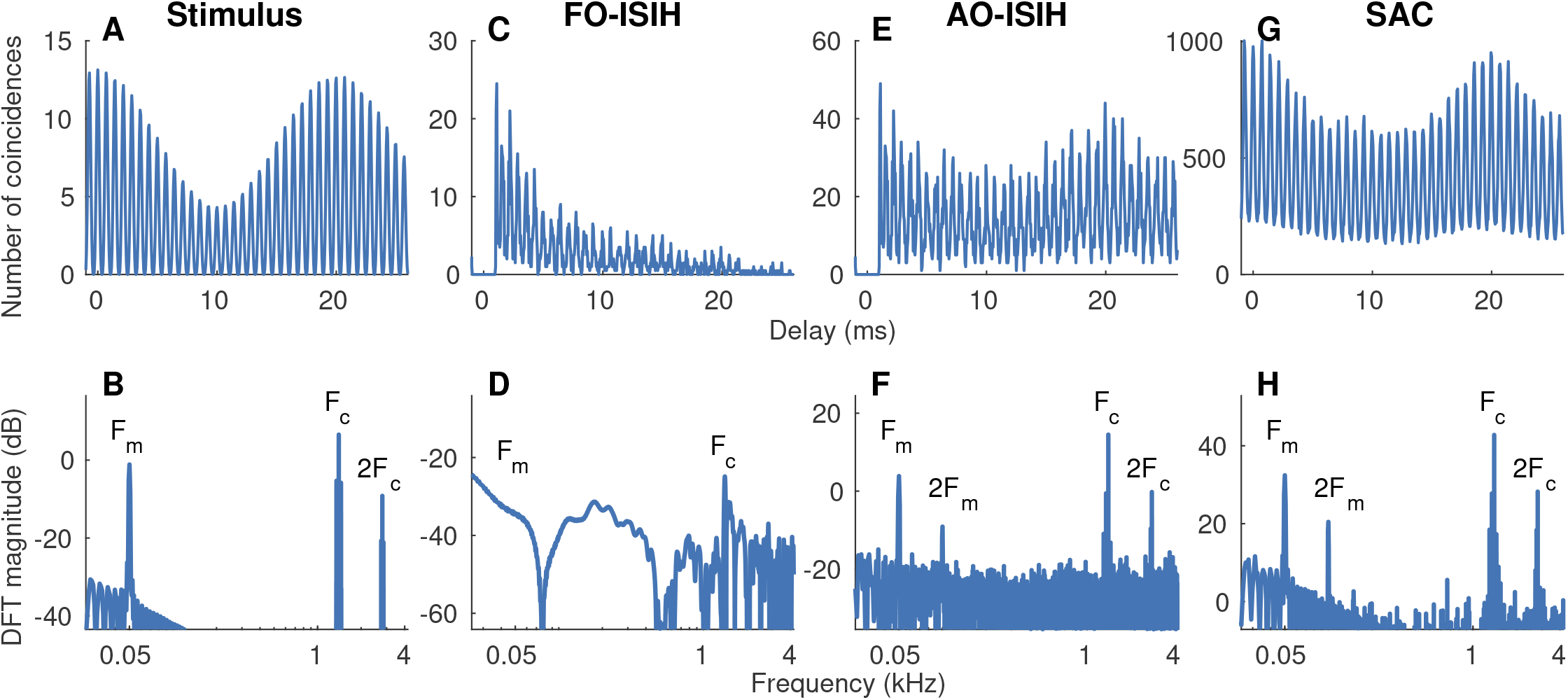
The shuffled autocorrelogram is better than the first-order and all-order ISI histogram, both in the time and frequency domains. Example correlograms (top) and associated spectra (bottom) constructed using spike trains recorded from an AN fiber (CF = 1.4 kHz, medium SR) in response to a SAM-tone at *F*_*c*_ = CF (50-Hz modulation frequency or *F*_*m*_, 0-dB modulation depth, 700-ms duration, 27 repetitions, 50 dB SPL). (A) Autocorrelation function of the half-wave rectified stimulus. (B) The discrete Fourier transform (DFT) of A. (C) The first-order (FO) ISI histogram. (D) DFT of C. The first-order ISI histogram poorly captures the carrier (TFS) and fails to capture the modulator (ENV). (E) The all-order (AO) ISI histogram. (F) DFT of E. The all-order ISI histogram captures both the carrier and modulator despite being noisy. Both the first-order (C) and the all-order (E) ISI histograms show dips for intervals less than the refractory period (∼0.6 ms), with the corresponding spectra corrupted by these refractory effects. (G) The shuffled autocorrelogram. (H) DFT of G. The shuffled autocorrelogram is smoother compared to the other correlograms, which also leads to improved SNR in the spectrum at both the carrier and modulator frequencies. All these ISI histograms are corrupted by rectifier distortion at twice the carrier frequency (2*F*_*c*_). Bin width = 50 *μ*s for histograms in C, E, and G.

**S1 Fig.**
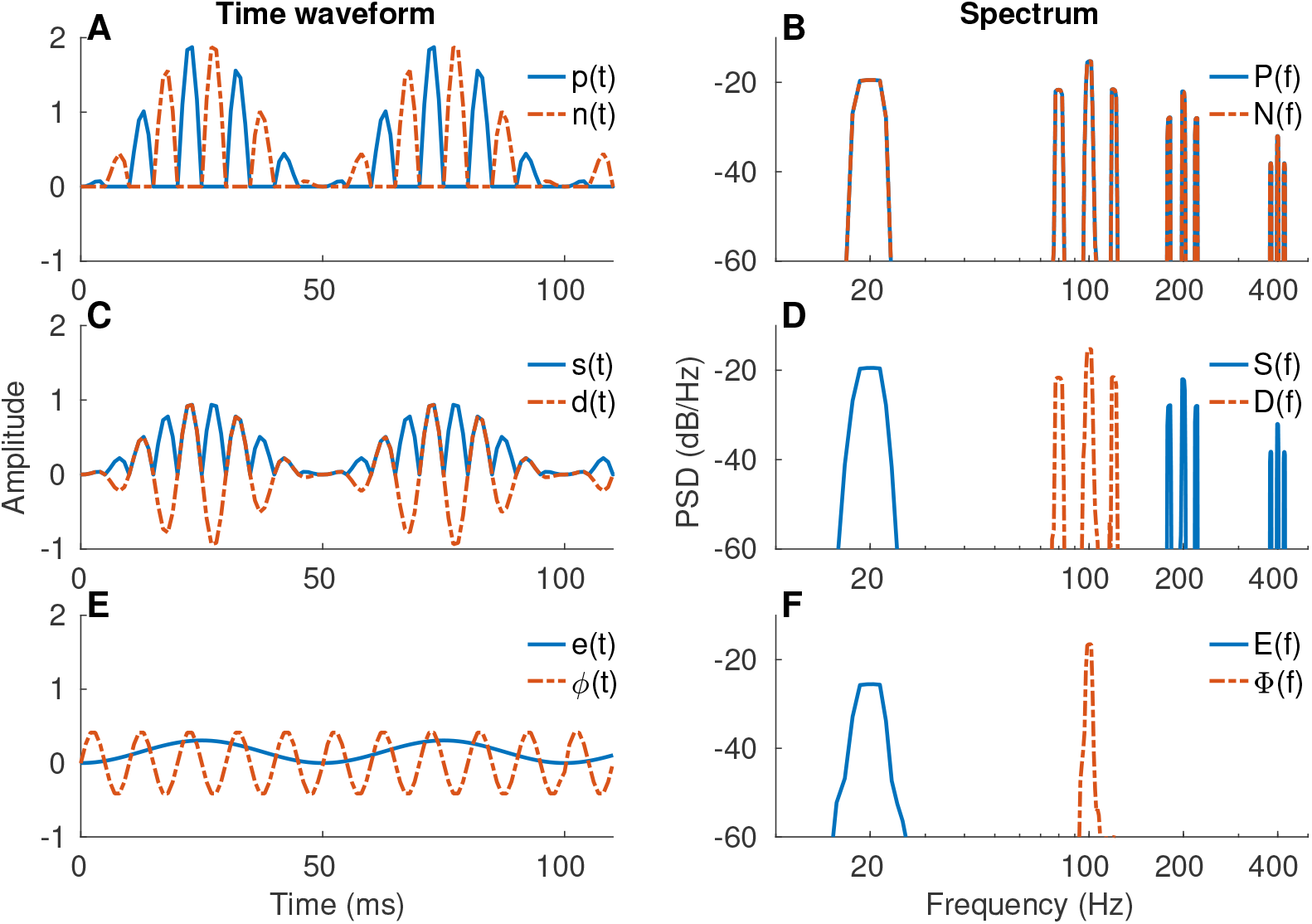
Graphical illustration of *apPSTHs* in Table 1. Graphical illustration for several *apPSTHs* for a simple half-wave rectifying model. A SAM tone (carrier = 100 Hz, modulation frequency = 20 Hz, sampling frequency = 1 kHz, duration = 1 s) was used as the stimulus, although for clarity only the first 100-ms are shown in time. Note that rectifier distortions occur at even harmonics of the carrier for *P* (*f*), *N* (*f*), and *S*(*f*), but not for *D*(*f*), *E*(*f*), or Φ(*f*).

**S2 Fig.**
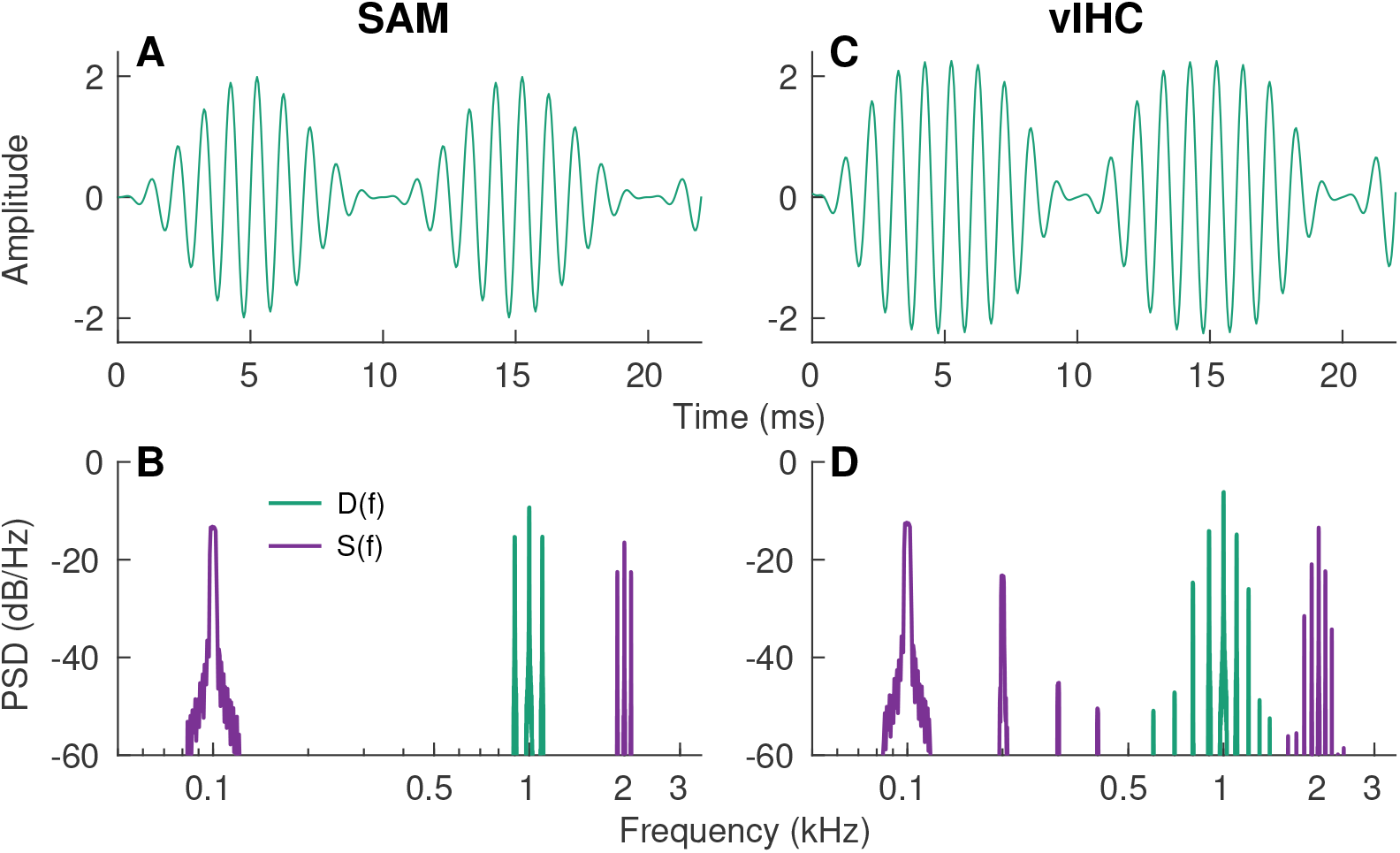
Nonlinear inner-hair-cell transduction function introduces additional sidebands in the spectrum for a SAM tone. (A) Waveform for a SAM tone (*F*_*c*_=1 kHz, *F*_*m*_=100 Hz, 0-dB modulation depth). (B) *D*(*f*) and *S*(*f*) for the SAM tone in A. (C) Waveform of the output after processing the SAM tone through a sigmoid function. The sigmoid function was used as a simple proxy for the inner-hair-cell transduction function. This output (vIHC) was further low-pass filtered at 2 kHz to mimic the membrane properties of inner hair cells. (D) *D*(*f*) and *S*(*f*) for the signal in C. In addition to having power at *F*_*c*_ and *F*_*c*_ ± *F*_*m*_, *D*(*f*) for vIHC has substantial energy at *F*_*c*_ *±* 2*F*_*m*_ (plus reduced energy at higher multiple *F*_*m*_-offsets from *F*_*c*_). Similarly, *S*(*f*) for vIHC has substantial energy at *F*_*m*_ as well as at the first few harmonics of *F*_*m*_. *S*(*f*) is also corrupted by rectifier distortion at 2*F*_*c*_ (and multiple *F*_*m*_-offsets from 2*F*_*c*_) as expected.

**S3 Fig.**
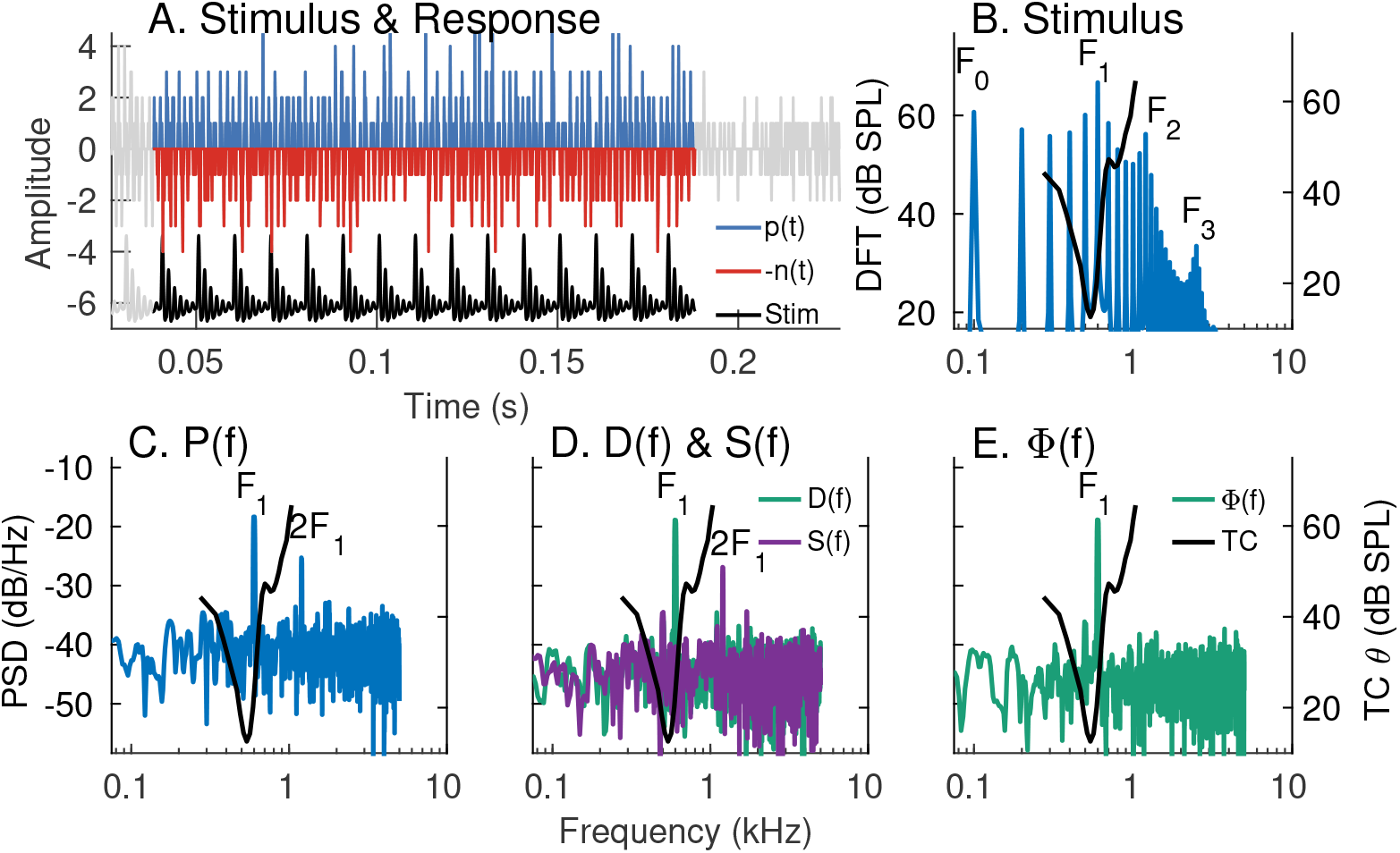
Neural characterization of ENV and TFS using *apPSTHs* for a synthesized stationary vowel Fig S3. Spectral-domain application of various *apPSTHs* to spike trains recorded in response to a stationary vowel. Example of spectral analyses of spike trains recorded from an AN fiber (CF= 530 Hz, SR=90 spikes/s) in response to a synthesized stationary vowel (*s*_1_ described in *Materials and Methods*, fundamental frequency: *F*_0_ = 100 Hz, first formant: *F*_1_ = 600 Hz). (A) Time-domain representation of *p*(*t*), *n*(*t*), and the stimulus (*Stim*). *n*(*t*) is flipped along the y-axis for display. Signals outside the analysis window are shown in gray. PSTH bin width = 0.1 ms. Number of stimulus repetitions per polarity = 30. Stimulus intensity = 65 dB SPL. (B) Stimulus spectrum (blue, left yaxis). In panels B-E, the frequency-threshold tuning curve (TC *θ*, black) of the neuron is plotted on the right y-axis. The neuron’s CF was close to the first stimulus formant. (C) *P* (*f*), which shows a strong response to the 6th harmonic (first formant) and the 12th harmonics (due to rectifier distortion). (D) Spectra for difference [*D*(*f*), green] and sum [*S*(*f*), purple] PSTHs. *D*(*f*) shows a clear peak at the 6th harmonic and little energy near the 12th harmonic. Similar to *P* (*f*), *S*(*f*) shows substantial energy at twice the TFS (*F*_1_) frequency due to rectifier distortion. (E) Spectra of Hilbert-based TFS PSTH [Φ(*f*), green]. *P* (*f*) and *S*(*f*) are corrupted by rectifier distortion at 2*F*_1_ frequency. The response primarily reflects TFS-based *F*_1_ coding (E) and little envelope coding (D), which is consistent with the “synchrony-capture” phenomenon for stationary vowel coding (Delgutte and Kiang, 1984a; Young and Sachs, 1979). Note that *E*(*f*) is not shown because *e*(*t*) was essentially flat across the vowel duration, and therefore had little energy other than at 0 Hz.

**S4 Fig.**
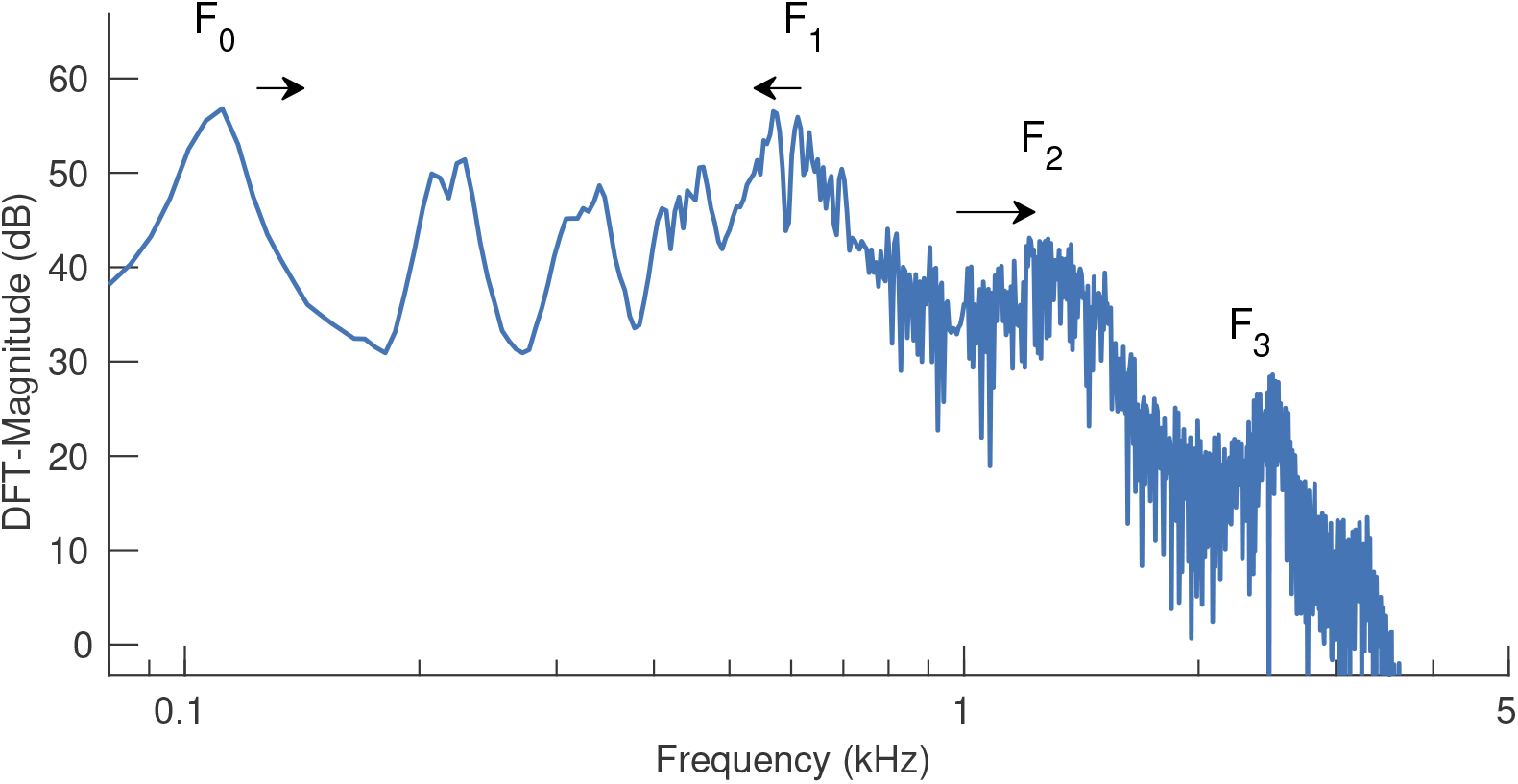
DFT-magnitude for the nonstationary vowel, *s*_2_. The stimulus duration was 188 ms. The movements of *F*_0_ (100 to 120 Hz), *F*_1_ (630 to 570 Hz), and *F*_2_ (1200 to 1500 Hz) are indicated by arrows. *F*_3_ was fixed at 2500 Hz.

**S5 Fig.**
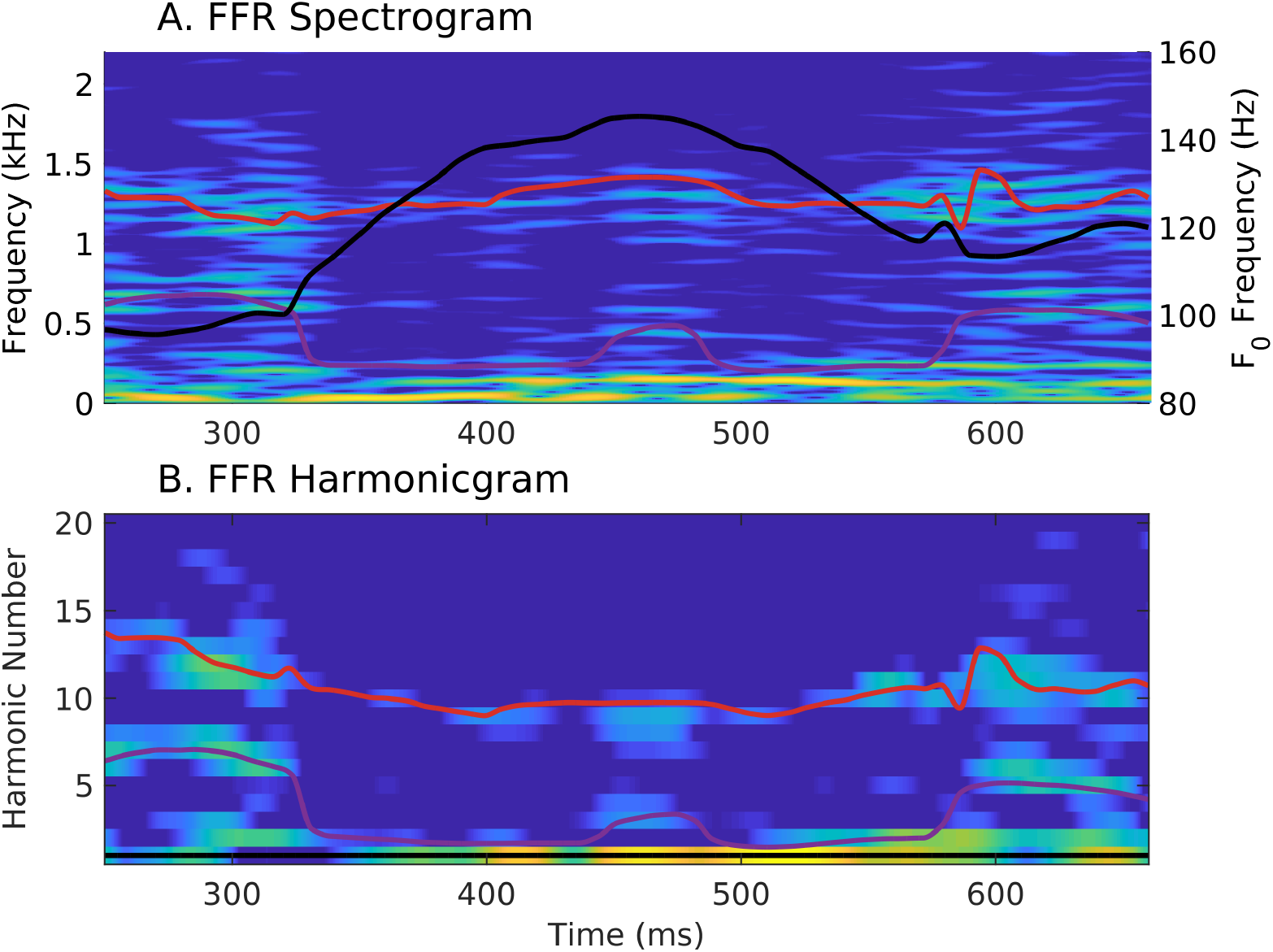
FFR harmonicgram can be constructed using the Hilbert-phase response. Same format as Fig 12. The spectrogram (A) and the harmonicgram (B) were constructed using *ϕ*(*t*).

